# Molecular mechanism of M1-mediated influenza A virus assembly

**DOI:** 10.64898/2026.04.26.720559

**Authors:** Hui Guo, Iosune Ibiricu, Margot Riggi, Robin Mohr, Raffaella Villa, Tillman Schäfer, Zhe Liu, Marius Boicu, John A. G. Briggs

## Abstract

Influenza A virus assembly is orchestrated by matrix protein 1 (M1), which engages ribonucleoproteins, mediates glycoprotein incorporation, and scaffolds lipid envelopment, but how these activities are performed and coordinated is unclear. We determined high-resolution structures of M1 within virions, virus-like particles, and in vitro. We found that M1 binds phosphatidylinositol 4,5-bisphosphate at the membrane, oligomerizes through an electrostatic interface, and switches between three alternative conformational states in different regions of the virion. The M1 conformer at the front associates with ribonucleoproteins and initiates budding. In the body of the virion, two conformers alternate to create an extended filament with seam-like discontinuities. Enrichment of the seam-prone conformation at the virion rear induces envelope closure and exposes a binding site for the cytoplasmic tail of neuraminidase (NA) to promote virus release. Conformational switching within a polarized M1 lattice thereby couples organization of the viral components with shaping of the virion architecture.

## Introduction

Influenza A virus (IAV) drives annual seasonal epidemics and poses an ongoing pandemic threat. The virion is pleiomorphic – often filamentous when isolated from infected hosts, but commonly bacilliform after lab adaptation (*1*, *2*). The genome consists of eight single-stranded, negative-sense, RNA segments. Each segment is bound by nucleoprotein (NP) and the polymerase complex to form the viral ribonucleoproteins (vRNPs) that localize towards the front end of filamentous virions in a characteristic “7+1” arrangement (*3–5*). The virion is surrounded by a host-derived lipid bilayer containing the glycoproteins hemagglutinin (HA) and neuraminidase (NA), as well as the low-copy-number ion channel matrix protein 2 (M2). HA, broadly distributed across the virion surface (*3*, *6*), mediates receptor binding and membrane fusion (*7*). NA is less abundant and is often enriched at the rear tip of filamentous virions (*3*, *6*), where it cleaves sialic acids to promote viral release and prevent self-aggregation. The inner surface of the lipid bilayer is coated by matrix protein 1 (M1), the most abundant and most conserved viral protein (*3*, *8*).

Assembly of influenza components occurs at the apical plasma membrane (*9*), particularly within lipid raft domains enriched in cholesterol and sphingomyelin (*10*, *11*). To reach budding sites, HA, NA, and M2 traffic via the ER–Golgi secretory pathway (*12*), vRNPs are transported to the cell periphery on Rab11-positive membranes (*13*, *14*), while recruitment of M1 from the cytoplasm is proposed to involve viral glycoproteins (*15*, *16*) and M2 (*17*, *18*).

M1 functions as the central organizer of virus assembly. It engages negatively charged lipids (*19–21*) and assembles into a polarized helix beneath the apical plasma membrane to induce extrusion of the filamentous virus particle. During assembly, M1 coordinates particle formation through interactions with vRNPs (*3*, *8*, *22*, *23*) and supports glycoprotein incorporation, a process influenced by the cytoplasmic tails of HA and NA (*15*, *16*, *24–26*). After entry, M1 disassembles in response to endosomal acidification to release vRNPs (*27*, *28*).

Structurally, M1 consists of 252 residues organized into N-terminal (NTD) and C-terminal (CTD) domains. The membrane-associating NTD has a compact, helical fold and has been observed in multiple crystal packing arrangements (*29*, *30*). Low-resolution cryo-electron tomography (cryo-ET) of intact virions shows that the CTD, which is largely unstructured in solution, becomes ordered and interacts with neighboring NTDs to promote M1 oligomerization (*8*). Even in the absence of membranes, purified recombinant M1 can assemble into distinct helical oligomers in vitro (*8*, *31*). However, how these assemblies relate to the native matrix in virions has not been established.

Currently, there is a lack of mechanistic and structural understanding of virus particle assembly: how are the front, the body, and the rear of the particle constructed; how are the glycoproteins and vRNPs engaged and positioned to coordinate assembly while establishing virus front-rear directionality; and how does M1 play its central role in these processes?

## Results and discussion

### M1 exhibits structural plasticity determined by two conformational switches

To provide the initial basis for understanding M1 structure and interactions, we first characterised assemblies formed by purified, recombinant M1 protein. Purified M1 protein is known to assemble into three different filamentous, helical assemblies: a high-salt assembly of the V97K mutant (*31*), nucleotide-associated assembly formed under alkaline conditions (*8*), and a high-salt assembly of wild-type (WT) M1 (*31*). Structures of the V97K and nucleotide-associated assemblies were previously determined at 3.4 Å and 3.8 Å resolutions, respectively.

We determined high-resolution cryo-electron microscopy (cryo-EM) structures of all three assemblies (2.9 Å, 2.1 Å, and 2.8 Å) using helical reconstruction followed by symmetry expansion and focused refinement of a local lattice patch (Fig. 1, A to C, and figs. S1 to S3).

**Fig. 1.**
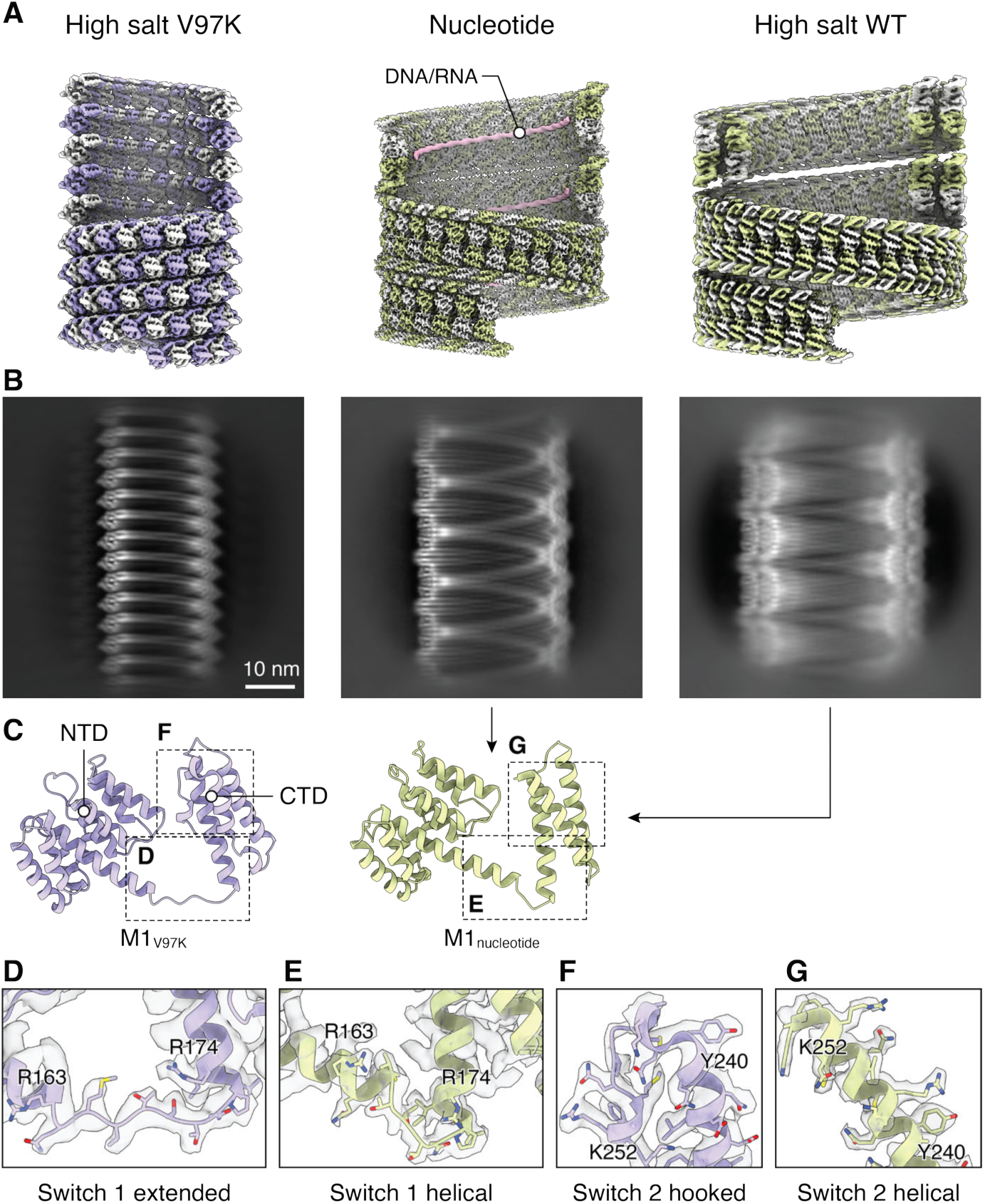
Structures of in vitro M1 assemblies reveal two conformational switches. (**A**) Helical reconstructions of V97K, nucleotide-associated and high-salt WT in vitro assemblies of M1, shown as surface views with alternate M1 molecules in color and in white for clarity. (**B**) Representative 2D class averages of the assemblies. Scale bar 10 nm. (**C**) Atomic models of the M1 monomer from the assemblies. The nucleotide-associated and WT high-salt assemblies adopt the same M1 conformation, and one representative model is shown. Boxes indicate the switch regions highlighted in (D to G). (**D–E**) Switch 1 (R163–P171 linker) is extended in M1_V97K_ and partially helical in M1_nucleotide_. (**F–G**) Switch 2 (C-terminal residues Y240–K252) forms a helix-loop-3_10_ helix (“hooked”) structure in M1_V97K_ and is helical in M1_nucleotide_.

Multilayered M1 assemblies are observed in influenza A virus-infected cells and the layers have been shown to resemble the nucleotide-associated assemblies (*32*, *23*). We found that the WT high-salt assembly has the same monomer structure and helical arrangement as the nucleotide-associated assemblies but is multilayered. We performed cryo-focused ion beam (cryo-FIB) milling of A/Hong Kong/1/1968 (H3N2, HK68) infected cells, and confirmed by two-dimensional (2D) classification that the WT high-salt assembly structure is representative of the multilayered M1 assemblies within the nucleus of infected cells (fig. S4).

Comparison of the M1 monomer structures in the three assemblies revealed two conformational switches in M1. Switch 1 involves residues R163-P171 in the NTD-CTD linker region: they form an extended linker in the V97K assembly (M1_V97K_) (Fig. 1D), but become partially helical in the monomer state shared by the nucleotide-associated and WT high-salt assemblies (M1_nucleotide_) (Fig. 1E and fig. S5A). Switch 2 primarily involves the C-terminal residues Y240–K252, but is associated with broader structural changes in the CTD: in M1_V97K_ these residues fold back to form a hooked configuration with a short 3_10_ helix (Fig. 1F), whereas in M1_nucleotide_ they extend the final alpha helix of M1 (Fig. 1G and fig. S5B). We therefore asked: how is the structural flexibility in M1 that is provided by these two conformational switches relevant for virus assembly?

### The virus body contains two, distinct M1 conformations

To address this, we aimed to derive the high-resolution structure of M1 within influenza virus particles. To avoid purification artifacts, we performed single-particle cryo-EM on filamentous HK68 virions budding directly from infected cells (Fig. 2A). HK68 was observed budding both as individual virions, and as bundles of virions. As expected, HA coated most of the envelope surface, NA accumulated toward the rear tip, vRNPs concentrated at the front, and M1 formed a continuous layer beneath the membrane (Fig. 2B). As reported previously by cryo-ET, the M1 layer exhibits substantial heterogeneity in diameter and helical start number (*8*). Even within individual virions, the front part of the viral filament body, including the vRNP-containing region, is often wider than the longer, rear part of the body (fig. S6A). We therefore used iterative 2D classification to group M1 filament segments from the virion body, first by radius and then by helical start (fig. S7 and Fig. 2C). Helical refinement was performed on each class, and segments with different helical symmetries but similar local packing were merged to obtain two asymmetric reconstructions of the local M1 lattice within the virus at 3.0 Å resolution (figs. S6 and S7 and Fig. 2, D and E).

**Fig. 2.**
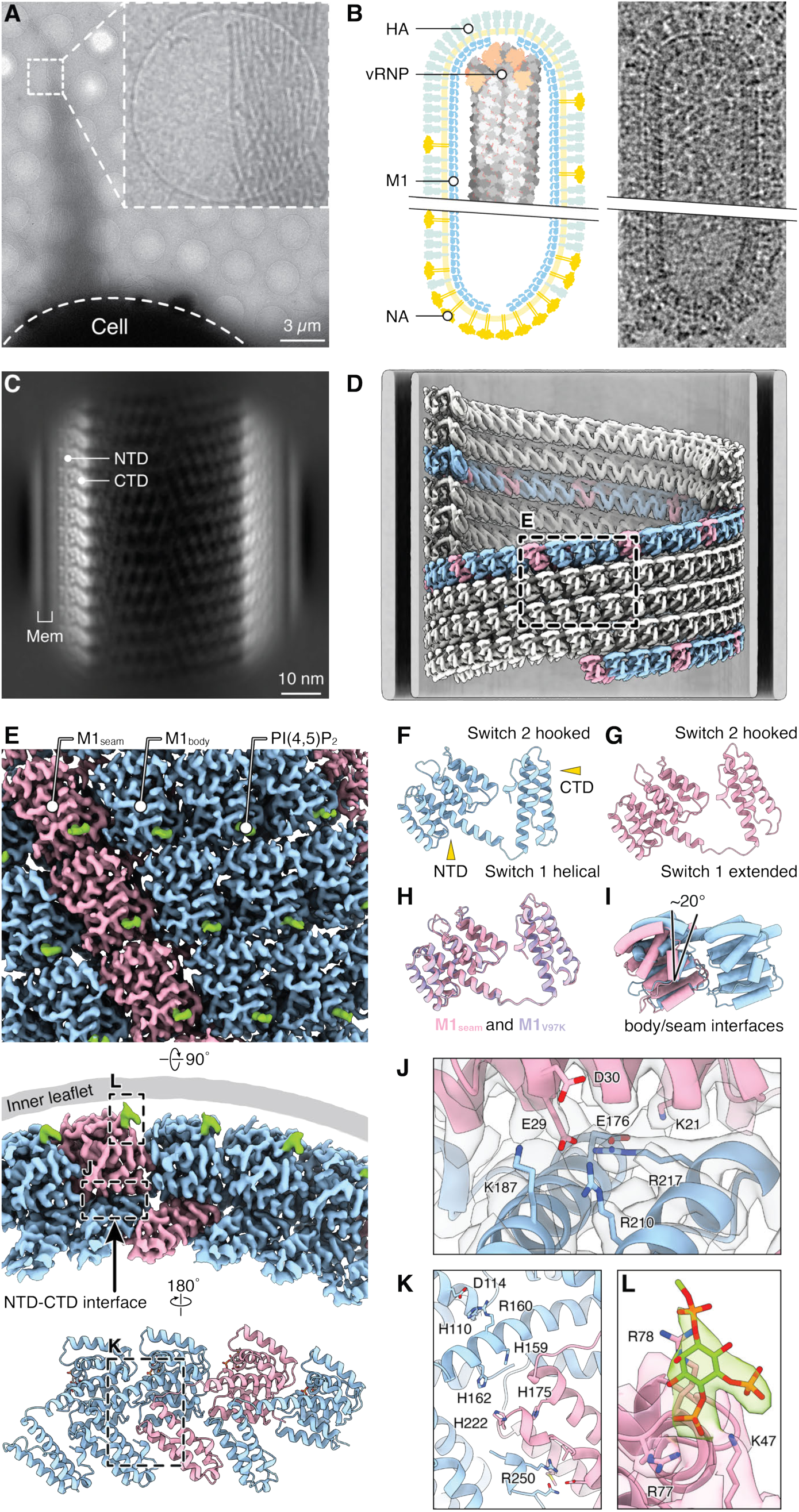
High-resolution in-virus structures of M1. (**A**) Representative micrograph of filamentous A/Hong Kong/1/1968 (H3N2, HK68) virions budding from infected cells (cell body is marked). Bundles of filaments are frequently observed (inset). Scale bar 3 µm. (**B**) Cartoon (left) and representative micrograph (right) illustrating virion organization, with proteins marked: HA coats the viral surface, NA is enriched toward the rear, vRNP density toward the front of the virion, and the M1 layer underlies the lipid bilayer. (**C**) Representative 2D class average of an HK68 virion segment from the majority, narrower population. The N and C-terminal domains (NTD and CTD) of M1 are marked, as is the lipid bilayer. Scale bar 10 nm. (**D**) Representative helical reconstruction of HK68 M1 with four helical starts. One helical strand is colored. Box indicates the region highlighted in (E). (**E**) Surface views of the reconstruction of the M1 lattice viewed from outside the virus (top panel) and from the side (middle panel). Lower panel shows a 180° rotated side view of the model. M1_seam_, M1_body_, and PI(4,5)P₂ densities are colored pink, blue, and green, respectively. (**F**) M1_body_ adopts the helical Switch 1 state seen in M1_nucleotide_ and the hooked Switch 2 state seen in M1_V97K_. (**G**) M1_seam_ adopts the extended Switch 1 and hooked Switch 2 states as seen in M1_V97K_. (**H**) Structural overlay of M1_seam_ and M1_V97K_ showing close similarity and both switches in the same states. (**I**) Cartoon structural overlay of M1_seam_ and M1_body_ showing the 20° relative NTD. (J) Close-up of NTD–CTD intersubunit contacts in the assembled lattice. Charged residues mediating the contact are marked. (**K**) Close-up of the CTD-mediated intersubunit contact involving R250. Positions of five histidines that may act in pH-sensitive disassembly are indicated. (**L**) Close-up of membrane-proximal density consistent with a PI(4,5)P₂ headgroup and electrostatic contacts formed with K47, R77, and R78.

A small subset of the segments (8%) adopted the expected helical assembly composed of a single M1 conformation, which we term M1_body_ (Fig. 2, D and E, blue, fig. S6F, and fig. S7, B and C). These segments were wider than the bulk population and corresponded to the front part of the virus body, including the vRNP-containing region (fig. S6A, blue segments). In M1_body_, switch 1 is in the partially helical conformation seen in M1_nucleotide_, while switch 2 is in the hooked conformation seen in the M1_V97K_. M1_body_ therefore represents a hybrid of the two in vitro structures (Fig. 2F and fig. S5C).

Most filament segments (92%) deviated from this organization and contained a second M1 conformation interspersed with M1_body_ at a 1:3 ratio (Fig. 2, D and E, pink, fig. S6G, and fig. S7, B and C). This subset of filaments represented the bulk population and corresponded to the long, rear part of the virus body (fig. S6A, pink segments). This second M1 conformation creates a discrete discontinuity in lattice packing, reminiscent of the microtubule seam, and we therefore term it M1_seam_. M1_seam_ shares the state of both switches with M1_V97K_, and their overall tertiary structures are highly similar (Fig. 2, G and H). Switch 1 is therefore in the extended state in M1_seam_, but in the partially helical state in M1_body_. Driven by this conformational difference in the inter-domain linker, M1_seam_ shows a ∼20° counterclockwise rotation of the NTD relative to M1_body_ (Fig. 2I). Viral matrix helices are generally assumed to consist of a single protein conformation. Our data show that, unexpectedly, the influenza virus body contains two interspersed conformations of M1 forming seam-like discontinuities.

### M1-M1 and M1-lipid interactions within the virus

The high-resolution structures of M1 within the virion allow us to interpret the protein-protein and protein-lipid contacts that mediate influenza assembly. Although the NTD–NTD interfaces vary within a virion due to the coexistence of M1_body_ and M1_seam_, the NTD–CTD contact is conserved and forms an electrostatic network involving residues K21, E176, R217, E29, D30, K187, and R210 (Fig. 2J). This interface may provide a major stabilizing contribution to M1 oligomerization, with additional support from CTD-CTD contacts involving R250 (Fig. 2K). Previous work has suggested that five histidines in M1 may contribute to a pH sensitive disassembly switch (*8*, *31*). We see that H159, H162, H175, and H222 form a histidine cluster (Fig. 2K), while H110 lies at an M1–M1 interface, where it packs against a nearby charged pair (R160 and D114), a configuration commonly observed in pH-sensing histidines (*33–35*).

At the membrane interface, each M1 monomer binds a lipid headgroup density with a shape consistent with phosphatidylinositol 4,5-bisphosphate (PI(4,5)P₂) that forms electrostatic contacts with K47, R77, and R78 (Fig. 2L). The well-defined binding site suggests that M1 makes a specific PI(4,5)P₂ interaction that would help anchor M1 selectively at the apical plasma membrane, which is the site of influenza budding and the major cellular reservoir of PI(4,5)P₂ (*9*, *36*). During entry, endosomal acidification would weaken such electrostatic M1-lipid contacts, as proposed for other filamentous enveloped viruses (*37*, *38*), and together with protonation of the histidine cluster, lead to destabilization and disassembly of the matrix layer.

### Formation of the seam is an inherent property of M1 assembly

We next probed whether the switching of M1 conformation that leads to seam formation is an inherent property of M1, or whether it is triggered or controlled by additional viral components. We determined the M1 lattice structure in virus-like particles (VLPs) where the composition of viral proteins can be defined (Fig. 3, A and B, and figs. S8 and S9). VLPs were produced in HEK293T cells by expressing HA and M segment (HAM) or NA and M segment (NAM). Seams were present in both VLP types, indicating that seam formation does not require the full complement of viral proteins and is not uniquely dependent on a specific glycoprotein, consistent with it being an inherent property of M1.

**Fig. 3.**
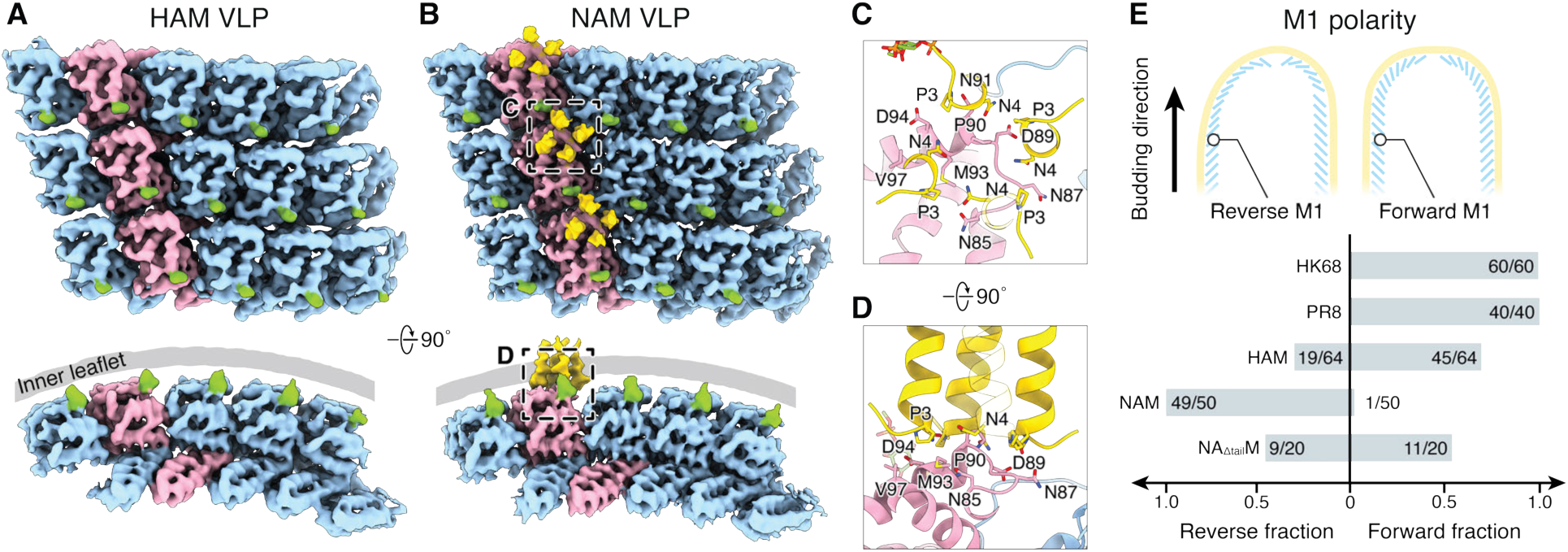
M1seam creates a binding platform for the NA tail. (**A**) Surface view of the unsharpened cryo-EM map of M1 from HAM VLPs (HA+M1+M2) showing the presence of both M1_seam_ (pink) and M1_body_ (blue) conformations. (**B**) Surface view of the unsharpened cryo-EM map of NAM VLPs (NA+M1+M2) showing four symmetrically arranged densities (yellow) between the membrane and M1_seam_ interface, corresponding to the N-terminal region of NA, including its cytoplasmic tail. (**C-D**) Model for the interaction between NA cytoplasmic tail and M1_seam_ viewed from outside the virus (C) and from the side (D). An AlphaFold3-predicted model of the NA N-terminal tetramer is refined into the density. P3 and N4 in the NA tails contact a hydrophobic patch in M1. (**E**) Quantification of M1 polarity across intact virions and VLPs. HK68 and PR8 virions show a consistent forward polarity, whereas HAM VLPs show mixed polarity. NAM VLPs show predominantly reverse polarity, but mixed polarity is seen in NAM VLPs where the NA cytoplasmic tail is deleted (NA_Δtail_M).

### M1_seam_ creates a binding platform for the NA tail

Although the tails of the glycoproteins have been implicated in virion assembly (*15*, *16*, *24*, *25*) and lie in close proximity to the M1 layer at the cytoplasmic face of the viral membrane, it has remained unknown if and how they interact with M1. In HAM VLPs we were unable to detect any density corresponding to the cytoplasmic tail of HA, suggesting that HA does not make any ordered interaction with M1. In NAM VLPs, however, we observed four symmetrically arranged densities associated with the membrane-binding face of M1_seam_ (Fig. 3B, yellow). An AlphaFold3-predicted model of the N-terminal region of NA shows that the highly-conserved cytosolic tail (MNPNQK) of the NA tetramer fits the densities well (fig. S10A). At this position, the rotated orientation of M1_seam_ repositions neighboring subunits to create a hydrophobic pocket, where residues P3 and N4 of the NA cytoplasmic tail contact N87, P90, N91 and M93 of the M1_seam_ (Fig. 3, C and D). At the corresponding position in M1_body_, the site is blocked by a positively charged surface from an adjacent M1, preventing NA binding (fig. S10, B and C). These data provide structural evidence for an NA–M1 interaction within the virion where the M1_seam_ conformation forms the NA binding platform.

### Consistent M1 polarity in virions requires other viral components

Influenza virions have clear directionality, with vRNPs at the front tip which buds first, and NA enriched toward the rear tip where scission and release occur (Fig. 2B). The underlying M1 lattice also has a defined polarity in which all CTDs point in one direction along the filament body (Fig. 2C) (*39*). We quantified the polarity of M1 relative to the directionality of budding and found that all observed virions had the same “forward” polarity in which the M1 CTD points away from the front tip, towards the rear tip (Fig. 3E and fig. S11, A and B). To test whether this polarity is intrinsic to M1 or whether it is imposed by other viral components, we examined the polarity of M1 in HAM and NAM VLPs. HAM VLPs displayed a mixture of forward and reverse polarity (Fig. 3E and fig. S11C), indicating that HA and M1 are not sufficient to impose a consistent M1 polarity in the absence of the other viral components. NAM VLPs displayed predominantly reverse polarity, where the M1 CTD points towards the front tip, instead of away from the front tip as in virions (Fig. 3E). Deleting the NA cytoplasmic tail (residues 2–6, NPNQK) in NAM VLPs eliminated this bias and restored mixed polarity (Fig. 3E and fig. S11D). These observations imply that the vRNP, or other viral components, are required to define the “forward” M1 polarity in virions. They further suggest that interactions between the NA tail and M1 may promote formation of a rear-tip-like organisation that, in the absence of the other components, can initiate filament formation with reverse M1 polarity.

### M1_nucleotide_ and M1_seam_ mediate formation of the front and rear viral tips

Superimposing the M1 arrangement from the virus body with the M1_nucleotide_ and M1_V97K_ in vitro assemblies revealed that these polymers adopt distinct curvatures (Fig. 4A, left), thereby repositioning the membrane-binding surface relative to the polar M1 filament axis (Fig. 4A, right). This relationship suggested to us that the M1_nucleotide_ conformation might be responsible for coating the virion front tip, whereas the M1_seam_/M1_V97K_-like conformation might form the rear tip—regions that differ in curvature and glycoprotein composition. We further noted that M1_nucleotide_ and M1_V97K_ are able to form substantially thinner filaments (285 Å and 250 Å, respectively (fig. S4D and fig. S1C) than the in-virus lattice (∼370-630 Å (fig. S7C)), as would be required for compatibility with the narrowing radius at the tips.

**Fig. 4.**
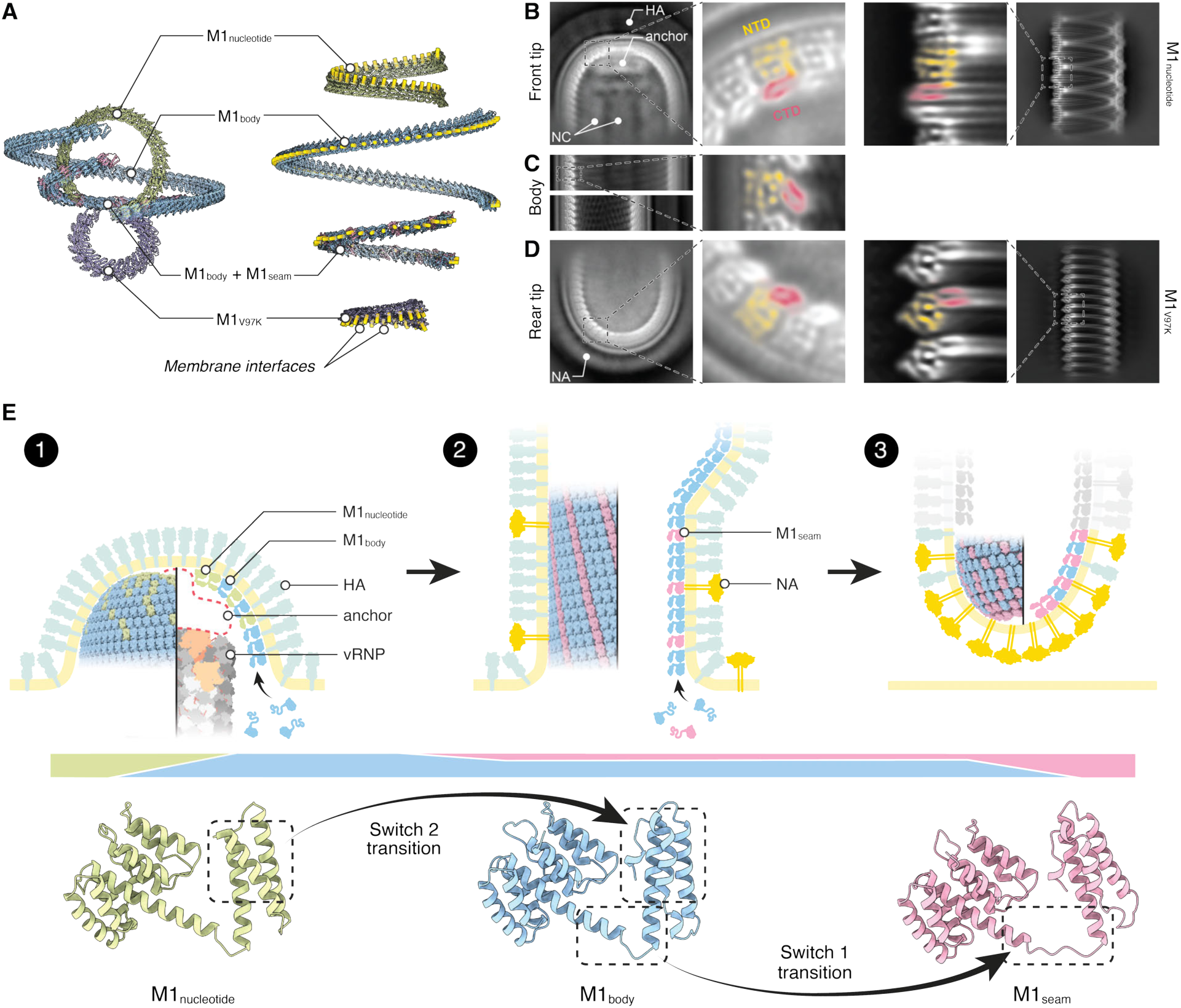
M1 conformations perform location-specific functions in virions. (**A**) Comparison of polymer propagation directions for M1_nucleotide_, M1_body_ and the mixed M1_body_-M1_seam_ strands seen in virions, and M1_V97K_. Polymers are aligned on their first monomer in the left panel and on the filament axis in the right panel. The membrane-interaction surfaces are highlighted by yellow bars – they point in different directions relative to the filament axis. (**B**) Left: representative 2D class average of HK68 front tips with a zoomed-in view of the boxed region. The HA, nucleocapsid (NC) and the anchor densities are marked. Right: representative 2D class average of M1_nucleotide_ with a zoomed-in view of the boxed region. The similar M1 orientation is indicated by the direction of the M1 CTD density (red), which points toward the outside of the filament, placing the membrane-binding M1 NTD density (yellow) towards the front of the filament. (**C**) Representative 2D class averages from the wider front part and the longer, narrower rear part of the HK68 filament body, with a zoomed-in view showing the CTD density (red) oriented downward relative to the filament axis. (**D**) Left: representative 2D class average of PR8 rear tips with a zoomed-in view of the boxed region. Right: representative 2D class average of the M1_V97K_ with a zoomed-in view of the boxed region. The similar M1 orientation is indicated by the direction of the CTD density (red), which points toward the inside of the filament, placing the membrane-binding M1 NTD density (yellow) towards the rear of the filament. (**E**) A model of M1-mediated influenza virus assembly. Step 1, at the front tip, M1_nucleotide_ (green) interacts with the nucleocapsid anchor and initiates polarized M1 assembly. The growing M1_nucleotide_ polymer incorporates increasing amounts of M1_body_ (blue) to shape the front tip of the virion. Step 2, M1_body_ polymerisation continues to extend the front of the virion, subsequently, M1_body_ and M1_seam_ (pink) polymerise at a 3:1 ratio to extend the long, filamentous rear body region. Step 3, progressive enrichment of M1_seam_ generates the rear tip of the virion. M1_seam_ provides a binding platform for the cytoplasmic tail of NA (gold), thereby enriching NA at the rear of the virion, where it fulfils its role in virus release. The prevalence of different states at different stages of assembly is illustrated by the coloured bar. The transition between M1_nucleotide_ and M1_body_ is mediated by switch 2 in the CTD. The transition between M1_body_ and M1_seam_ is mediated by switch 1 at the NTD-CTD linker. Two conformational switches thereby allow M1 to transition between three conformations within the assembling virus particle, changing the direction of polymerisation relative to the lipid bilayer and changing the interactions with other viral components.

To further test this model, we picked the tips of virus filaments from the HK68 dataset and collected cryo-EM data for the bacilliform strain A/Puerto Rico/8/1934 (H1N1, PR8) (fig. S12A). 2D classification of virion tips recapitulated the expected glycoprotein organization, with HA enriched at the front and NA at the rear in both strains (Fig. 4, B and D, and fig. S12B).

HK68 NA appears ∼45 Å longer than PR8 NA, due to its extended hypervariable stalk. The nucleocapsids are clearly visible in the 2D class averages. We observed additional ordered density layers within the first turn of M1 and extending toward the nucleocapsids. This structure may represent multiple copies of nuclear export protein (NEP) and the polymerase complex components of the vRNPs, but the composition cannot be defined from our data. Due to its position between M1 and the nucleocapsid, we term it the nucleocapsid anchor (Fig. 4B and fig. S12B).

The NTD and CTD of M1 are clearly visible adjacent to the membrane in the class averages (Fig. 4, B to D, yellow and red). As predicted, the orientation of M1 at the front of the virus matches that of M1_nucleotide_, while the orientation at the rear matches that of the M1_V97K_/M1_seam_ (Fig. 4, B and D), indicating that the two in vitro conformations correspond to tip-specific M1 states in virions. These tip-specific M1 states colocalize with specific viral components: M1_nucleotide_ and the genome segments are present at the front of the virion, where components of the nucleocapsid anchor may couple M1_nucleotide_ polymerisation with vRNP recruitment to define the forward M1 polarity seen in virions. M1_seam_ and NA are present at the rear of the virion, where the NA tail interacts with M1_seam_.

The NA cytoplasmic tail is highly conserved across IAV (*40*, *41*) and has been proposed to mediate M1 trafficking to the membrane (28, 35), promote NA incorporation (29, 36), and influence virion morphology (36, 37). However, NA lacking the cytoplasmic tail supports formation of filamentous NAM VLPs containing both M1 and NA (fig. S11D), indicating that the tail is not absolutely required for particle budding or NA incorporation, although it may enhance these processes. Our identification of M1_seam_ as an NA-tail-binding platform and our observation that the M1_seam_ coats the rear tip of the virion, together assign a function to the conserved NA tail and provide the structural mechanism explaining how NA is enriched at the rear end of the virion to promote virus release. This mechanism could act by a high concentration of the NA tail recruiting and stabilizing M1_seam_ to promote closure of the rear of the virion, or equivalently by a M1_seam_-rich virion rear forming and then capturing NA. These options are not mutually exclusive. Indeed, the previous observation that M1 can polymerise into the cytosol at the rear of budding particles (*23*) indicates that a M1_seam_-rich virion rear can form first, while our observation that the NA-M1_seam_ interaction is sufficiently strong to bias budding to reverse polarity in the absence of other viral components indicates that a higher concentration of NA tail can stabilize M1_seam_ to promote formation of a rear-tip architecture. Beyond its enrichment at the rear tip, M1_seam_ is also distributed along the filament body, creating additional NA-tail binding pockets that may facilitate particle separation and release in bundle-forming strains such as HK68.

We speculated that forming the intermediate curvatures connecting the virion body and tips could be facilitated by mixing M1_body_, M1_seam_, and M1_nucleotide_ conformations in varying proportions, aided by flexibility inherent within each conformation. To test this idea, we constructed geometric computational models (see Materials and Methods) in which the fraction of M1_seam_ increases toward the rear and the fraction of M1_nucleotide_ increases toward the front: models built with these mixtures were able to continuously coat tip-like geometries (figs. S13 and S14).

### A model for influenza virus assembly

Although structures of individual influenza proteins are well characterized, how they are coordinated to build a directional, infectious particle has remained unclear. The data presented here frame a mechanism for virus particle assembly (Fig. 4E). M1 engages the apical plasma membrane via PI(4,5)P₂ and forms a linear polymer through an electrostatic NTD-CTD interface. Two conformational switches allow M1 to adopt three distinct monomer structures within the assembling virus particle, each of which polymerises in a different direction relative to the lipid bilayer. This provides a basis for the M1 conformation to be biased by the local membrane geometry, and vice versa. At the front tip, M1_nucleotide_ interacts with the nucleocapsid anchor and initiates polarized M1 assembly. The growing M1_nucleotide_ polymer incorporates increasing amounts of M1_body_ to shape the front tip of the virion. The body elongates around the vRNPs via polymerisation of M1_body_ alone, later narrowing to form a long-extended filament made of M1_body_ and M1_seam_ at a 3:1 ratio. Toward the rear, progressive enrichment of M1_seam_ generates the rear-tip architecture. M1_seam_ provides a binding platform for the cytoplasmic tail of NA, thereby enriching NA at the rear of the virion, where it fulfils its essential role in virus release. In this way, conformational switching allows polymerising M1 to shape the front, body, and rear of the filamentous virion, and to directly couple membrane shaping to the respective front and rear localisation of the vRNPs and NA. During entry, endosomal acidification weakens PI(4,5)P₂–M1 interactions and activates histidine-based pH sensors within the matrix lattice, destabilizing the M1 layer and triggering its disassembly.

## Materials and methods

### Cell lines

Human embryonic kidney 293T (HEK293T; CRL-3216), and Madin–Darby canine kidney (MDCK; CCL-34) cells were obtained from the American Type Culture Collection (ATCC) and were cultured in Dulbecco’s modified Eagle medium (DMEM) GlutaMAX (Gibco) supplemented with 10% fetal bovine serum (FBS) and 1% penicillin–streptomycin. HEK293T cells were used to produce influenza VLPs. MDCK cells were used to propagate HK68 and PR8 viruses for single-particle cryo-EM studies and for cryo-FIB preparation of HK68-infected cells. All experiments with viruses were carried out under BSL-2 conditions. All incubations were performed at 37 °C and 5% CO₂ unless otherwise noted.

### Preparation of HK68 and PR8 virions on grids for cryo-EM without purification

Influenza A/Hong Kong/1/1968 (H3N2, HK68) and Influenza A/Puerto Rico/8/1934 (H1N1, PR8) starting stocks, generated by reverse genetics (*23*), were obtained from Dr. Petr Chlanda (Heidelberg University). Viruses were expanded in MDCK cells as described previously (*8*). For on-grid production of virions, Quantifoil AU-200 mesh R2/1 holey carbon grids (Quantifoil Micro Tools) were glow-discharged and pre-equilibrated for 1 h in 35 mm × 10 mm dishes containing 2 mL cell culture medium. MDCK cells were seeded onto grids at 0.5–1 × 10^5^ cells per dish and incubated for ∼20 h to allow cell attachment to the grids. Prior to infection, dishes were washed with DMEM lacking FBS and supplemented with 0.3% BSA. Cells on grids were inoculated with diluted HK68 or PR8 at a multiplicity of infection of 3 and incubated for 1 h with gentle mixing every 15 min. The inoculum was removed and replaced with DMEM containing 0.3% BSA and L-(tosylamido-2-phenyl) ethyl chloromethyl ketone (TPCK)-treated trypsin at 0.5–5 µg/mL, adjusted based on trypsin activity. Grids were plunge-frozen ∼48 h post-infection. HK68-infected grids were used for single-particle cryo-EM to determine the M1 structure in virions, for analysis of M1 layer polarity in HK68, and for cryo-FIB milling to prepare lamellae of HK68-infected cells. PR8-infected grids were used for cryo-ET to analyze M1 layer polarity in PR8.

### Purification of PR8 virus for 2D classification of virion tips

PR8 infections were performed as for on-grid preparation, except that 6 × 10^6^ MDCK cells were seeded in a T175 flask. Culture supernatant was collected 48 h post-infection and supplemented with 2 mM MgCl₂ and benzonase at 300 U/mL. The supernatant was clarified by centrifugation at 2,000 × g for 10 min at room temperature, then layered onto a 30% (w/v) sucrose cushion prepared in centrifugation buffer (10 mM HEPES, 100 mM NaCl, 1 mM EDTA, pH 7.4). Virus was pelleted by ultracentrifugation at 26,000 rpm for 90 min at 4 °C using an SW32 Ti rotor in an Optima L-90K ultracentrifuge (Beckman). The pellet was resuspended in Dulbecco’s phosphate-buffered saline (DPBS) overnight, dialyzed against DPBS using a 25 nm MCE membrane (MF-Millipore) for 2 h at 4 °C, and used within 1 day.

### Preparation of VLPs on grids for cryo-EM without purification

HA, NA, and the M segment from HK68, as well as NA from Influenza A/Singapore/1/57 (H2N2, SG57) and its cytosolic-tail deletion variant, NA_Δtail_ (residues 2–6 removed), were cloned into the pCAGGS backbone (35). Quantifoil AU-200 mesh R2/1 holey carbon grids (Quantifoil Micro Tools) were glow-discharged and placed in 35 mm × 10 mm dishes containing 2 mL medium for 1 h. HEK293T cells were seeded onto grids at 1.0–1.5 × 10^5^ cells per dish and incubated for ∼20 h. VLPs were generated by FuGENE-mediated transfection (Promega) following the manufacturer’s instructions. HAM VLPs were produced by co-transfecting HK68 HA and HK68 M at a 1:2 ratio. HK68 NA together with HK68 M did not produce filamentous VLPs under our conditions, so NAM VLPs were produced using SG57 NA with HK68 M at a 1:2 ratio. NA_Δtail_M VLPs were produced using SG57 NA_Δtail_ with HK68 M at a 1:2 ratio. Grids were plunge-frozen ∼48 h post-transfection.

### Cryo-EM specimen preparation

Cryo-EM specimens of influenza A virus and VLPs produced on grids were prepared similarly. Grids containing cells with virus or VLPs were removed from the culture medium, and 2 µL of medium was added immediately prior to plunge-freezing into a 1:1 ethane/propane mixture using a Leica EM GP2 at 100% humidity and 20 °C with a blot time of 5 s. For purified PR8 virus, 3 µL of diluted virus was applied to glow-discharged Quantifoil Cu-200 mesh R2/1 holey carbon grids (Quantifoil Micro Tools) and plunge-frozen into a 1:1 ethane/propane mixture using a Leica EM GP2 at 100% humidity and 4 °C with a blot time of 5 s.

### Lamella preparation of HK68-infected cells by cryo-FIB milling

Plunge-frozen HK68-infected MDCK cells on grids were clipped into cartridges, mounted in a shuttle with 45° pretilt, and transferred to an Aquilos 2 cryo-FIB (Thermo Fisher Scientific). The sample surface was coated with organometallic platinum using the gas injection system three times for 30 s each. The stage was tilted to 15° for milling. Rough milling was performed above and below the cell of interest using gallium ions at 0.5–0.1 nA to generate lamellae of ∼400 nm thickness. Lamellae were then polished at 50–30 pA to a final thickness of 150–200 nm. The FIB acceleration voltage was 30 kV for all milling steps. Milling progress was monitored using the scanning electron microscope operated at 3 kV and 13 pA.

### Cryo-EM data acquisition for on-grid preparations of HK68 virus and HAM and NAM VLPs

Grids were imaged with a Titan Krios G4 transmission electron microscope operated at 300 kV and equipped with a cold field emission gun, a Falcon4i direct detector, and a Selectris X energy filter (ThermoFisher Scientific). Unpurified virus and VLP preparations yielded few particle-containing holes, about five holes per grid square on average. Low magnification maps of all grid squares were acquired at a nominal magnification of 580× using a 4 s exposure (total dose, 0.14 e^−^/nm^2^), large defocus (−500 µm), and an inserted energy filter to improve visibility of filamentous particles. Holes containing virus or VLPs were then identified manually. Higher-magnification images of the selected holes were collected at 11,500×, and final acquisition positions were manually marked on each hole image. Final data collection used stage moves to the selected holes followed by micrograph acquisition using beam-image shift, scripted in SerialEM (*42*). For HK68 virus, one dataset was collected targeting the filament body. Because virions tended to cluster together, tips were frequently captured within the same fields of view. For HAM and NAM VLPs, two datasets were collected for each VLP. One targeted the filament body for M1 structure determination, and the other targeted filament tips for 2D classification.

All datasets were collected at 130,000×, giving a calibrated pixel size of 0.9534 Å. Data were recorded in electron-event representation (EER) format (*43*). The exposure rate was 8 to 9 e^−^/pixel/s and the total exposure was 40-50 e^−^/Å^2^. Detailed acquisition parameters and the number of micrographs for each dataset are provided in tables S1 to S3.

### Cryo-EM data acquisition for purified PR8 virus

Purified PR8 virus grids were imaged on the same Titan Krios G4 microscope with identical hardware as described above. Data collection was performed using EPU (ThermoFisher Scientific) instead of the SerialEM-based workflow. Data were recorded in EER format at 130,000× nominal magnification under exposure conditions similar to the previous datasets.

### Cryo-ET data acquisition and tomogram reconstruction for unpurified PR8 virus

Grids containing MDCK cells infected with PR8 virus were imaged on the same Titan Krios G4 microscope with identical hardware as described above. Ten budding sites where PR8 virions remained attached to the budding membrane were identified, and tilt series were acquired from −60° to 60° in 3° increments using a dose-symmetric tilt scheme in Tomography (ThermoFisher Scientific). Data were recorded in EER format at 105,000× nominal magnification, corresponding to a calibrated pixel size of 1.2142 Å. The exposure time per tilt was 0.5 s with a dose of 3 e^−^/Å^2^ per tilt, resulting in a total dose of 123 e^−^/Å2 per tilt series. Tomograms were reconstructed using AreTomo3 (37) and subsequently denoised with cryoCARE (38) prior to analysis of M1 layer polarity in budding PR8 virions (fig. S11B).

### Cryo-EM and cryo-ET data acquisition and tomogram reconstruction for M1 filaments in lamellae of HK68-infected cells

Grids containing cryo-FIB milled lamellae of HK68-infected cells were imaged on the same Titan Krios G4 microscope with identical hardware as described above. Multi-layered M1 filaments were observed predominantly in the nucleus of infected cells (fig. S4A). Tilt series and single-exposure micrographs were collected using Tomography (ThermoFisher Scientific). Tilt series acquisition and processing were performed as described for unpurified PR8 virus (fig. S4B). Micrographs were collected under the same optical and detector settings as the tilt series, but as single exposures with a total dose of 53 e^−^/Å².

### M1 layer polarity analysis in virus and VLPs

For HK68 virus, filament tips with clear vRNP density were assigned as the front budding tip. M1 layer polarity was determined by examining the orientation of the M1 CTD (Fig. 4, B to D, red). Forward polarity was assigned when the M1 CTD pointed away from the budding direction (fig. S11A). Reverse polarity was assigned when the M1 CTD pointed in the same direction as budding. PR8 virions are much shorter, and vRNP density is often close to both tips, which makes budding direction difficult to assign from vRNP alone. We therefore analyzed tomograms of PR8 virions that remained attached to the budding membrane. Budding direction was defined by the membrane attachment, and M1 polarity was surveyed by examining the CTD orientation (fig. S11B). For VLPs, vRNP density is absent and cannot be used to define budding direction. We therefore analyzed micrographs of VLPs that retained a small remnant of budding membrane. M1 polarity was estimated from the CTD orientation relative to the inferred VLP growth direction (fig. S11, C and D).

### Cryo-EM image processing of M1 in HK68 virus

The data processing workflow is summarized in fig. S7. Unless noted otherwise, all image processing was performed in CryoSPARC v4 (*44*). From 4,335 micrographs, 514,558 helical segments were picked using crYOLO v1.9.2 (*45*) with a dataset-specific trained model. Particles were extracted with a box size of 960 pixels and Fourier-cropped to 256 pixels for 2D classification. About 8% of particles formed three 2D classes with clearer internal M1 features and larger diameters, which were later identified as seamless filaments. The remaining particles formed 2D classes with less well-resolved internal M1 features, consistent with seam-containing filaments (fig. S7B). Each subset was subjected to one additional round of 2D classification, performed separately for seam-containing and seamless particles. M1 layer diameters were measured from the resulting 2D class averages as the distance across the outer edges of the M1 density layer, and summarized as a diameter distribution (fig. S7C).

For high-resolution reconstruction of seam-containing filaments, segments with M1 layer diameters of 384 to 412 Å were re-extracted with a box size of 864 pixels and Fourier-cropped to 256 pixels, then split into four diameter groups. Within each group, additional 2D classification was performed to separate classes with different helical starts (fig. S7D). Helical parameters were estimated for each 2D class by measuring the pitch from the 2D class average, followed by trial-and-error searches to determine the number of subunits per turn and the number of helical starts. Helical refinement was performed separately for each 2D class. In total, 60 2D classes were refined, and the resulting reconstructions revealed that multiple classes shared the same helical parameters. Classes with matching helical parameters were then combined, yielding 15 distinct helical parameter sets. Eleven parameter sets with three to five helical starts were selected for further processing. For these, particles were re-extracted at a box size of 864 pixels without binning and processed with one CTF-refinement cycle consisting of helical refinement, CTF refinement, and a final round of helical refinement to obtain the final maps for each parameter set (fig. S7E).

Although these maps had different helical parameters, the local M1 packing was similar and could be combined. Symmetry expansion was performed using the helical parameters of each class, followed by signal subtraction of a 3 × 4 M1 patch. Subtracted particles were real-space cropped to a box size of 256 pixels and locally refined, yielding a 3.3 Å map. Further 3D classification in RELION-5.0 (*46*) separated particles into four seam-containing classes with different seam positions and two classes without a seam. The four seam-containing classes corresponded to neighboring, near-equivalent M1 registers, so one class was selected for further refinement. Iterative CTF refinement and Blush regularization (*47*) in RELION-5.0 yielded a 3.1 Å map. Further 3D classification and refinement produced a final reconstruction at 3.0 Å resolution from 129,614 particle images (fig. S7F). Gold-standard Fourier shell correlation (FSC) curves for this and all subsequent final reconstructions are shown in fig. S15.

For the seamless M1 reconstruction, helical refinement was initiated from one seamless 2D class and processed using a similar workflow, yielding a final reconstruction at 3.0 Å resolution from 265,061 particle images (fig. S7G).

### Cryo-EM image processing of M1 in NAM and HAM VLPs

The overall processing workflows for M1 in VLPs were similar to those used for HK68 virus (figs. S8 and S9), with the differences described below. NAM VLP M1 layers were more homogeneous than those in virions (fig. S9, B and C). Most helical segments could be classified into 10 classes spanning a range of diameters, with either +1 or −1 helical start. For high-resolution reconstruction, three classes with similar helical parameters were combined, yielding a 3.3 Å resolution map from 91,384 particle images (fig. S9D). Signal subtraction was repeated using the HK68 mask to enable direct map comparison.

HAM VLPs showed heterogeneity more similar to the HK68 virus dataset (fig. S8B). For simplicity, we selected one class and applied a similar workflow, resulting in a 3.5 Å map from 79,234 particle images (fig. S8C).

### Cryo-EM image processing of virus and VLP tips

To obtain 2D classes of virus and VLP tips (Fig. 4, B and D, and fig. S12B), dataset-specific crYOLO models were trained to pick filament tips. For HK68 on-grid, purified PR8, and the tip-targeted HAM VLP and NAM VLP datasets, 4,106, 14,284, 4,334, and 2,359 particles were extracted with a box size of 864 pixels and Fourier-cropped to 216 pixels prior to 2D classification.

### Cryo-EM image processing of M1 filaments in lamellae of HK68-infected cells

To obtain 2D classes of M1 filaments in infected cells (fig. S4C), multi-layered M1 filaments were manually picked in RELION-5.0 as filaments and imported into CryoSPARC v4. In total, 7,669 helical segments were extracted with a box size of 1024 pixels and Fourier-cropped to 256 pixels prior to 2D classification. To obtain top-view classes, 7,538 particles were manually selected, extracted with a box size of 512 pixels, and Fourier-cropped to 128 pixels prior to 2D classification. The diameter of the innermost M1 layer was estimated by fitting concentric circles to the M1 layers (fig. S4D).

### M1 Construct expression

Full-length M1 from PR8, including WT and V97K, was cloned into pET-21. The WT construct carried a C-terminal LEHHHHHH extension. Protein was expressed in *E. coli* Rosetta™ 2(DE3) using auto-induction medium at 24 °C for 24 h (*48*). Cells were harvested by centrifugation at 4000 × g for 15 min.

### WT M1 purification and high-salt tube assembly

Approximately 4 g of cell pellet were resuspended in 50 mL Ni-NTA buffer (50 mM sodium phosphate, 500 mM NaCl, 10% v/v glycerol, 10 mM imidazole, 1 mM TCEP, 2 M urea, pH 8.0) supplemented with 2 mM MgCl₂, *S. marcescens* nuclease (300 U/mL; produced in-house by the Max Planck Institute of Biochemistry Protein Production Core Facility), and protease inhibitor cocktail (1 tablet; [Roche]). Cells were lysed with two passes through an EmulsiFlex-C5 homogenizer (Avestin) at 15,000 to 20,000 psi. Cell debris was removed by centrifugation at 20,500 rpm at 4 °C for 30 min using a JA-25.50 rotor (Beckman Coulter). The supernatant was filtered through a 0.45 µm filter and loaded onto a 1 mL GoBio Mini Ni-NTA column (Bio-Works) equilibrated in Ni-NTA buffer. Bound M1 was washed with Ni-NTA buffer supplemented with 5 mM ATP, and 10 mM MgCl₂, and eluted with Ni-NTA buffer containing 250 mM imidazole. The eluate was exchanged into SP buffer (50 mM sodium citrate, 150 mM NaCl, 1 mM TCEP, 1 M urea, pH 5.0) using a PD-10 desalting column and loaded onto a HiTrap SP HP 1 mL column (Cytiva) equilibrated in SP buffer. M1 was eluted using a linear NaCl gradient, with M1-containing fractions eluting at approximately 450 mM NaCl. Fractions containing M1 were pooled and stored at −80 °C. To assemble M1 into tubes, purified M1 was diluted to 0.1 mg/mL in 200 mM HEPES and 2 M NaCl and incubated at 4 °C for 24 h. For cryo-EM specimen preparation, assembled M1 tubes were dialyzed against DPBS using a 25 nm MCE membrane (MF-Millipore) for 2 h at 4 °C.

### WT M1 nucleotide-associated assembly purification

Nucleotide-associated M1 filaments were previously produced by mixing purified M1 with plasmid DNA (*8*). In our hands, WT M1 expressed in *E. coli* formed assemblies closely resembling the published nucleotide-associated filaments, although with one fewer nucleotide strand. Purification followed the WT protocol described above, except that urea was omitted from all buffers and SP cation-exchange chromatography was not performed. After Ni-NTA purification, fractions containing M1 were pooled and stored at 4 °C until use.

### V97K M1 purification and high-salt tube assembly

Full-length M1 from PR8 carrying the V97K mutation was cloned into pET-21. Purification was performed as described previously (*31*). Briefly, 11 g of cell pellet were resuspended in 100 mL size-exclusion chromatography (SEC) buffer (50 mM HEPES, pH 7.0, 150 mM NaCl, 2 M urea, 2 mM DTT) supplemented with *S. marcescens* nuclease (300 U/mL; produced in-house by the Max Planck Institute of Biochemistry Protein Production Core Facility). Cells were lysed by sonication for 3 min, and cell debris was removed by centrifugation at 10,000 × g at 4 °C for 30 min. RNA was precipitated by adding LiCl to 0.5 M final concentration and incubating at 4 °C for 30 min, followed by centrifugation at 10,000 × g at 4 °C for 30 min. Impurities were precipitated by addition of solid ammonium sulfate to 10% (w/v) final concentration and incubation at 4 °C for 30 min, followed by centrifugation at 10,000 × g at 4 °C for 30 min. M1 was then precipitated from the supernatant by increasing ammonium sulfate to 30% (w/v) final concentration by addition of solid ammonium sulfate and incubating at 4 °C for 30 min, followed by centrifugation at 10,000 × g at 4 °C for 30 min. The M1 pellet was dissolved in SEC buffer and further purified by size-exclusion chromatography on a Superdex 200 HiLoad 26/60 column (Cytiva) equilibrated in SEC buffer. Fractions containing M1 were pooled based on SDS-PAGE analysis and loaded onto a 5 mL HiTrap Q HP column and a 5 mL HiTrap SP HP column (Cytiva) connected in series and equilibrated in ion-exchange buffer (50 mM HEPES, pH 7.0, 150 mM NaCl, 1 M urea). After sample loading, the HiTrap Q HP column was disconnected, and M1 was eluted from the HiTrap SP HP column using a linear NaCl gradient from 150 mM to 1 M NaCl in ion-exchange buffer. M1 fractions were pooled and dialyzed into storage buffer (50 mM HEPES, pH 7.0, 1 M urea) and stored at −80 °C. To assemble V97K M1 into tubes, purified protein was diluted to 1.5 mg/ml in 3M NaCl, and incubated at 37 °C for 24 h.

### Cryo-EM specimen preparation and acquisition for M1 in vitro assemblies

All cryo-EM specimens of M1 in vitro assemblies were prepared similarly. A 3 µL volume of WT high-salt assemblies, WT nucleotide-associated assemblies, and V97K high-salt assemblies at 0.1 mg/mL, 0.2 mg/mL, and 1.5 mg/mL, respectively, was applied to glow-discharged Quantifoil Cu-200 mesh R2/1 holey carbon grids (Quantifoil Micro Tools). Grids were plunge-frozen into a 1:1 ethane/propane mixture using a Leica EM GP2 at 100% humidity and 4 °C with a blot time of 2 s. Grids containing M1 in vitro assemblies were imaged under the same conditions as purified PR8 virus. Data were collected using EPU (ThermoFisher Scientific) on the same Titan Krios G4 microscope with identical hardware, and were recorded in EER format at 130,000× nominal magnification with similar exposure conditions.

### Cryo-EM image processing of in vitro M1 tube assemblies

Processing followed the same overall strategy as for the virus and VLP datasets (figs. S5 to S7). Briefly, helical segments were extracted and subjected to 2D classification to select tube populations for helical refinement. Where applicable, particles were cleaned by 3D classification and transferred to RELION-5.0 for Bayesian polishing and Blush refinement. For combined reconstructions, symmetry expansion and signal subtraction were performed using the refined helical parameters, followed by local refinement.

V97K M1 high-salt tube assembly (fig. S1). From 9,444 micrographs, 2,787,465 helical segments were extracted with a box size of 448 pixels and Fourier-cropped to 144 pixels. 2D classification separated helical filaments from a minor population of stacked rings (fig. S1B). For the helical subset, 2,432,914 segments were subjected to consensus helical refinement, cleaned by 3D classification with helical symmetry in RELION-5.0, and refined to 2.7 Å from 224,598 segments. Symmetry expansion and signal subtraction using a mask containing six M1 subunits were then performed. Subtracted particles were real-space cropped to 256 pixels, followed by 3D classification and local refinement, yielding a final filament reconstruction at 2.9 Å resolution from 679,491 particle images. Stacked rings were processed separately by generating an initial model using helical reconstruction with a small rise, followed by local refinement with C21 symmetry. Duplicates arising from helical picking were removed prior to a final local refinement, yielding a 3.0 Å stacked-ring reconstruction from 90,618 particle images.

M1 nucleotide-associated assembly (fig. S2). From 7,091 micrographs, 715,290 helical segments were extracted with a box size of 640 pixels and Fourier-cropped to 320 pixels. Two rounds of 2D classification showed that most tubes formed filaments containing a single nucleotide on the inside, with a minor population of stacked M1 rings (fig. S2B). Filament classes were separated into two diameter groups of ∼330 Å and ∼338 Å. Segments were re-extracted with an unbinned 640-pixel box size and refined separately, yielding 2.4 Å maps for both groups after Bayesian polishing and final helical refinement. The two groups were then combined by symmetry expansion and signal subtraction using a mask containing six M1 subunits. Subtracted particles were real-space cropped to 256 pixels and locally refined, producing a final reconstruction at 2.1 Å resolution from 2,348,388 particle images.

WT M1 high-salt tube assembly (fig. S3). From 16,198 micrographs, 2,214,851 helical segments were extracted with a box size of 640 pixels and Fourier-cropped to 192 pixels. Class averages indicated reduced stability of multilayer tubes, so multiple rounds of 2D classification were used to remove damaged and heterogeneous particles. A subset of 287,684 two-layer segments was selected for reconstruction. Because the inner and outer layers had different helical symmetry, each layer was refined separately from the same segments by subtracting the other layer, yielding intermediate maps at 4.1 Å for the inner layer and 4.3 Å for the outer layer. For each layer, symmetry expansion and signal subtraction using a mask containing six M1 subunits were performed, followed by real-space cropping to 256 pixels, local refinement, 3D classification, and Blush refinement in RELION-5.0. Final reconstructions reached 2.8 Å for the inner layer from 210,893 particles and 2.9 Å for the outer layer from 155,491 particles.

### Atomic model building

Initial atomic models were generated using dataset-specific approaches. For the M1_nucleotide_ assembly, the initial model was built with ModelAngelo v1.0 (*49*), and this model was subsequently used as the starting model for refinement of both the M1_nucleotide_ and M1 WT high-salt assembly. For the M1_V97K_ assembly, PDB 7JM3 was used as the initial model. Initial models for M1 in virions with and without seams were independently generated with ModelAngelo v1.0; the virion-with-seam model was then used as the starting model for M1 in NAM VLPs. For the NA model in the NAM VLP structure, the N-terminal 20 residues of the NA tetramer were predicted with AlphaFold3 and used as an initial model. All models were manually adjusted in Coot-1 (*50*) and ISOLDE v1.10.1 (*51*), followed by real-space refinement in Phenix-2.0 (*52*). Model validation statistics are reported in table S1.

### Models of M1 polymers with different ratios of M1 conformations

Our data suggested that virion front and rear tips are enriched in M1_nucleotide_ and M1_seam_, respectively, whereas the filament body is formed by either M1_body_ alone or a mixed M1_body_:M1_seam_ lattice at a 3:1 ratio. To test whether the intermediate polymer curvatures needed to coat the tips could be generated by mixing M1 conformations, we built polymer models with varying ratios of M1_nucleotide_ and M1_body_, or of M1_body_ and M1_seam_, in UCSF ChimeraX (*53*) using *matchmaker* command (fig. S13, A and B). These models showed that mixed helices can generate a continuum of membrane-binding-surface orientations.

To analyse the polymerization direction and the orientation of the membrane-binding surface relative to the polymer, we defined a reference frame based on an M1_body_-only filament (fig. S7G), with the y axis aligned with the virus filament growth direction, the z axis directed radially outward (normal to the membrane surface), and the x axis orthogonal to both (fig. S13C). For each experimental helix, the rotation between adjacent monomers was measured in UCSF ChimeraX using the *measure rotation* command after aligning the first monomer to this reference frame. The resulting rotation matrices were converted to Euler angles (𝜃𝑥, 𝜃𝑦, 𝜃𝑧) using an extrinsic XYZ convention (fig. S13C, circles). Average Euler angles for the predicted, mixed helices were then calculated by combining the adjacent-monomer Euler angles from the experimental helices in the appropriate ratios (fig. S13C, squares).

The radius of the helical filament (fig. S13D) was calculated as 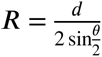, where 𝑑 is the distance between adjacent monomers and 𝜃 is the rotation angle between them. Values of 𝑑, measured in UCSF ChimeraX as NTD–NTD distances between consecutive monomers, were 28.87 Å for M1_body_, 28.74 Å for M1_nucleotide_, and 31.01 Å for M1_seam_. To determine the orientation of the membrane-binding surface (fig. S13E), one monomer from each predicted helix was aligned to the corresponding monomer in the M1_body_ reference helix, and the polymer axis was compared with that of the reference helix after projection into the yz plane.

### Models of M1 polymers at the front and rear tips of the virion

Step 1: Extraction of tip geometry

The shape of each tip was described from 2D class averages by manually tracing a smooth curve through the centres of the visible M1 NTD densities (fig. S14A, step 1). For each tip, the y axis was defined as the filament axis with 𝑦 = 0 at the point where the curved tip meets the straight body. The traced curve was represented in terms of radial distance 𝑅(𝑦) from the filament axis. The traced (𝑦, 𝑅) points were then fit with a weighted cubic smoothing spline to generate a continuous radial profile 𝑅(𝑦) for each tip (fig. S14A, step 1).

The local helical pitch, 𝑃 (𝑦), was estimated from the fitted radial profile 𝑅(𝑦). For each tip 2D class average, 𝑅(𝑦) passes through the centres of 𝑛 visible M1 layers. The arc length along 𝑅(𝑦) was therefore divided into 𝑛 − 1 equal segments (10 segments in the example shown in fig. S14A), defining successive turn points along the tip. 𝑃 (𝑦) was defined to be the 𝑦 distance between each pair of adjacent turn points at the 𝑦 value midway between the two turn points. 𝑃 (0) = 39.16 Å is defined as the pitch of the reference M1_body_ helix (fig. S7G). These points were then fit with a smooth spline to obtain a continuous pitch function, 𝑃 (𝑦) (fig. S14A, step 1). The orientation of the membrane-binding surface, 𝛼(𝑦), was derived from the slope of the fitted radial profile as 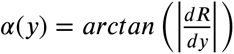, with 𝛼(0) = 0 enforced. Together, these procedures yielded three parameters describing each tip: radius 𝑅(𝑦), pitch 𝑃 (𝑦), and membrane-binding-surface orientation 𝛼(𝑦) (fig. S14A, step 1).

Step 2: Building a geometric model

Monomer coordinates were generated from the fitted functions 𝑅(𝑦) and 𝑃 (𝑦) by assuming a constant M1 NTD-NTD spacing 𝑑 along the underlying helical polymer. At each axial position 𝑦_𝑖_, the local radius 𝑅(𝑦_𝑖_) and pitch 𝑃 (𝑦_𝑖_) were used to calculate the helical path length per turn, 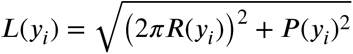. From this, the axial (𝛥𝑦_𝑖_) and azimuthal (𝛥𝜙_𝑖_) increments associated with one monomer step were obtained: 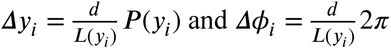. The monomer-center coordinates were then written as (𝑥_𝑖_, 𝑦_𝑖_, 𝑧_𝑖_) = (𝑅(𝑦_𝑖_) 𝑠𝑖𝑛 𝜙_𝑖_ , 𝑦_𝑖_, 𝑅(𝑦_𝑖_) 𝑐𝑜𝑠 𝜙_𝑖_) with iterative updates 𝑦_𝑖+1_ = 𝑦_𝑖_ + 𝛥𝑦_𝑖_ and 𝜙_𝑖+1_ = 𝜙_𝑖_ + 𝛥𝜙_𝑖_. For an N-start helix, the strands were assumed to be in phase, giving C_N_ symmetry. Each strand retained the same angular increment as the underlying helical sequence, while the axial rise per strand became 𝑁𝛥𝑦_𝑖_. Thus, for strand 𝑚, 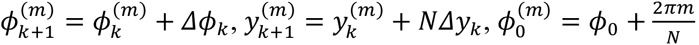, with 𝑚 = 0, … , 𝑁 − 1.

Monomer orientations were defined using a local reference frame based on the fitted tip geometry. In this frame, the local z-axis was oriented normal to the membrane-binding surface (fig. S14A, step 2, red arrows). The local x-axis was defined as the azimuthal tangent around the filament circumference (fig. S14A, step 2, blue arrows), and the local y-axis was defined as the cross product of z and x. This yielded an absolute orientation for each monomer along the tip without introducing additional twist beyond that required by the fitted geometry. Euler angles between consecutive monomers were then calculated from these absolute orientations using the convention defined above.

Step 3: Assignment of local filament composition and Euler angle adjustments

For each monomer in the geometric model, we measured its geodesic distance in Euler-angle space to each measured and predicted filament shown in fig. S13C, using the corresponding _(_𝜃_𝑥_, 𝜃_𝑦_, 𝜃_𝑧)_ values. Each monomer was then matched to the geodesically closest filament, thereby assigning a local filament composition. Consecutive monomers with the same predicted composition were grouped into segments, and conformations were assigned within each segment according to the corresponding ratio. For example, a segment matched to a 1:3 M1_nucleotide_:M1_body_ composition was modelled using one M1_nucleotide_ monomer and three M1_body_ monomers. The same approach was used for segments matched to a 1:3 M1_seam_:M1_body_ composition. In each case, the Euler angles from the experimental structures were used for the M1_nucleotide_ or M1_seam_ monomer, while the M1_body_ monomers were assigned a common set of Euler angles such that the segment-average Euler angles matched those of the geometric model.

We also considered an alternative model for front-tip formation in which there is no mixing of M1_nucleotide_ and M1_body_, but instead a single transition point (fig. S14C, right). In this model, each monomer in the geometric model was assigned as either M1_body_ or M1_nucleotide_ according to which experimentally observed conformation was geodesically closer in Euler-angle space, while the Euler angles of the geometric model were retained. This produced a single transition point in the polymer at which the conformation changes. As expected, the largest geodesic deviation from the M1_nucleotide_ angles was observed at the transition point (5.5° for HK68 virions and 5.7° for PR8 virions). As the in vitro assemblies are unlikely to sample the full M1 conformational space, we also considered a model allowing a small amount of additional inter-domain flexibility within the M1_nucleotide_ conformation. In this model, monomers assigned as M1_nucleotide_ were allowed an additional 2° of variability beyond the maximum deviation observed at the transition point in the previous model. Under this assumption, the transition point shifted to a different position along the polymer (fig. S14C, right, lower panel).

Step 4: 3D modelling

Molecular models of M1_body_, M1_nucleotide_, and M1_seam_ were first aligned to the reference frame defined above (fig. S13C). Surface representations of the models were then generated using the *molmap* command in UCSF ChimeraX. The resulting polygonal meshes were exported as .obj files, and imported into Autodesk Maya. Helical assemblies were constructed using 𝑁 independent hierarchical system(s) of parented mesh objects, with 𝑁 corresponding to the number of strands in the helix. Each object was assigned the coordinates calculated in step 2, together with the orientation and M1 conformation determined in step 3, with positions in “worldspace” set using the *xform* command and rotations set using the *setAttr* command.

## Acknowledgements

This work was funded by the Max Planck Society (J.A.G.B.) and the Deutsche Forschungsgemeinschaft (DFG, German Research Foundation), Projektnummer 240245660—SFB 1129 (J.A.G.B.). H.G. was supported by an EMBO Postdoctoral Fellowship (ALTF 306-2023). Cryo-EM data were collected in the Department of Cell and Virus Structure, Max Planck Institute of Biochemistry. We thank Dustin Morado for assistance with data collection; Florian Beck and Inga Wolf for assistance with computing infrastructure; Julia Peukes and Xiaoli Xiong for discussions of sample preparation; Aurelia Groessl for cloning; Petr Chlanda and Moritz Wachsmuth-Melm for sharing virus strains and for discussions on virus preparation; Petr Chlanda and Leo James for helpful comments on the manuscript; and Jürgen Plitzko for support of this work.

## Author contributions

H.G. and J.A.G.B. designed the research. I.I. prepared virus stocks and performed cryo-FIB milling. H.G. and I.I. prepared virus and VLP samples for cryo-EM and cryo-ET. R.M., R.V., M.B., and H.G. performed in vitro M1 experiments. H.G. collected cryo-EM and cryo-ET data with assistance from T.S. and Z.L. H.G. processed cryo-EM and cryo-ET data and built atomic models. M.R. and H.G. built virion tip models. H.G. and J.A.G.B. analyzed and interpreted data. H.G., M.R., and J.A.G.B. prepared the figures. H.G. and J.A.G.B. wrote the manuscript with input from all authors. J.A.G.B. obtained funding and managed the project.

## Data Availability

Structures determined by electron microscopy, including helical reconstructions and symmetry-expanded local patches, have been deposited in the Electron Microscopy Data Bank under accession codes EMD-57330 (M1_V97K_), EMD-57331 (M1_nucleotide_), EMD-57332 (M1 WT high-salt assembly), EMD-57326 (M1 in virions with seam), EMD-57327 (M1 in virions without seam), EMD-57328 (M1 in NAM VLP), and EMD-57329 (M1 in HAM VLP). Corresponding atomic models have been deposited in the Protein Data Bank under accession codes 29RJ (M1_V97K_), 29RK (M1_nucleotide_), 29RL (M1 WT high-salt assembly), 29RG (M1 in virions with seam), 29RH (M1 in virions without seam), and 29RI (M1 in NAM VLP).

---

**Fig. S1.**
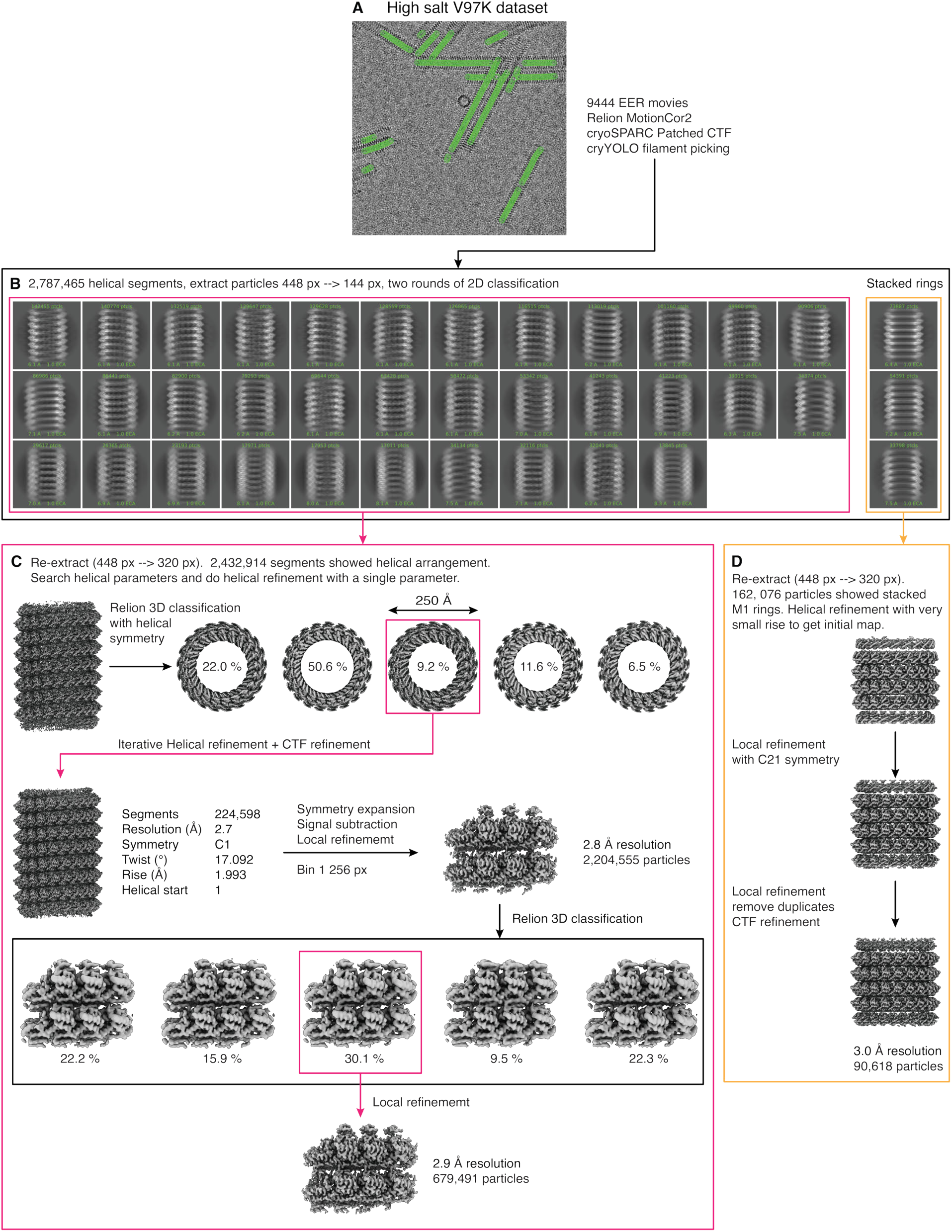
Image processing workflow for the M1V97K assembly. For a detailed description of the image processing, see materials and methods section. (**A**) Representative micrograph of the M1_V97K_ assembly with picked filament segments indicated (green). (**B**) Representative 2D class averages of the M1_V97K_ assembly. Most particles form helical filaments, while a minor population are tubes formed from stacked M1 rings. (**C**) Helical particles were subjected to a consensus helical refinement followed by 3D classification with helical symmetry to obtain a high-resolution helical reconstruction. Particles were then processed by symmetry expansion, signal subtraction using a mask including 6 M1 monomers, and iterative local refinement and 3D classification. M1_V97K_ tubes have an outer diameter of ∼250 Å, indicating that M1_V97K_ can assemble into tightly curved M1 polymers similar to those seen at virion tips (Fig. 4 and figs. S13 and S14). (**D**) For stacked-ring particles, an initial model was generated by treating the assembly as a helix with a small rise, followed by local refinement with C21 symmetry enforced. The final reconstruction was obtained after removing duplicate particles and performing a final local refinement.

**Fig. S2.**
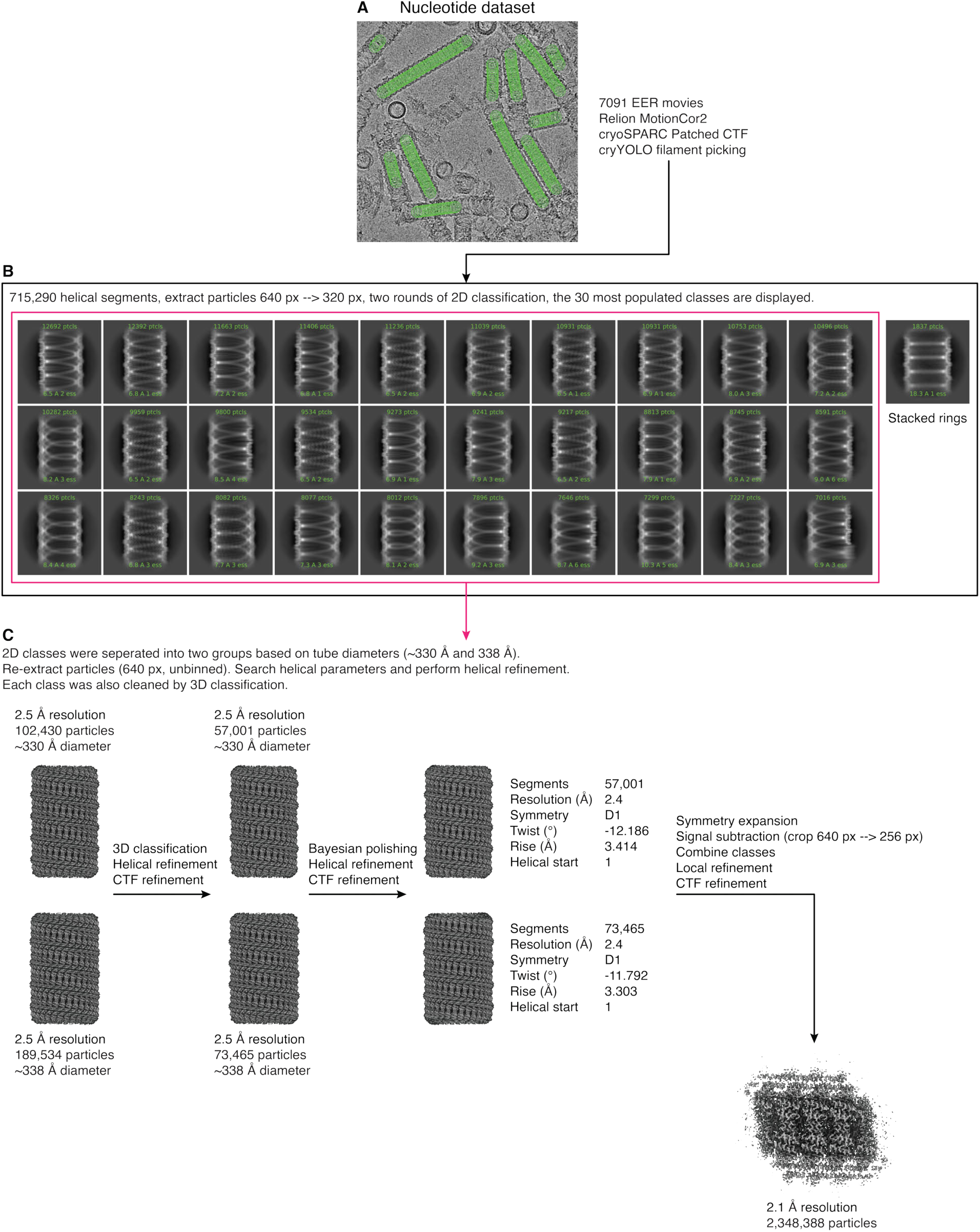
Image processing workflow for the M1nucleotide assembly. For a detailed description of the image processing, see materials and methods section. (**A**) Representative micrograph of the M1_nucleotide_ assembly with picked filament segments indicated (green). (**B**) Representative 2D class averages of M1_nucleotide_ assemblies. Most particles form filaments, whereas a minor population forms tubes with stacked M1 rings. (**C**) 2D classes were grouped into two subsets based on filament diameter. For each subset, helical parameters were searched and helical refinement was performed. The two subsets were then merged and processed by symmetry expansion, signal subtraction using a mask including 6 M1 monomers, and iterative local refinement and 3D classification.

**Fig. S3.**
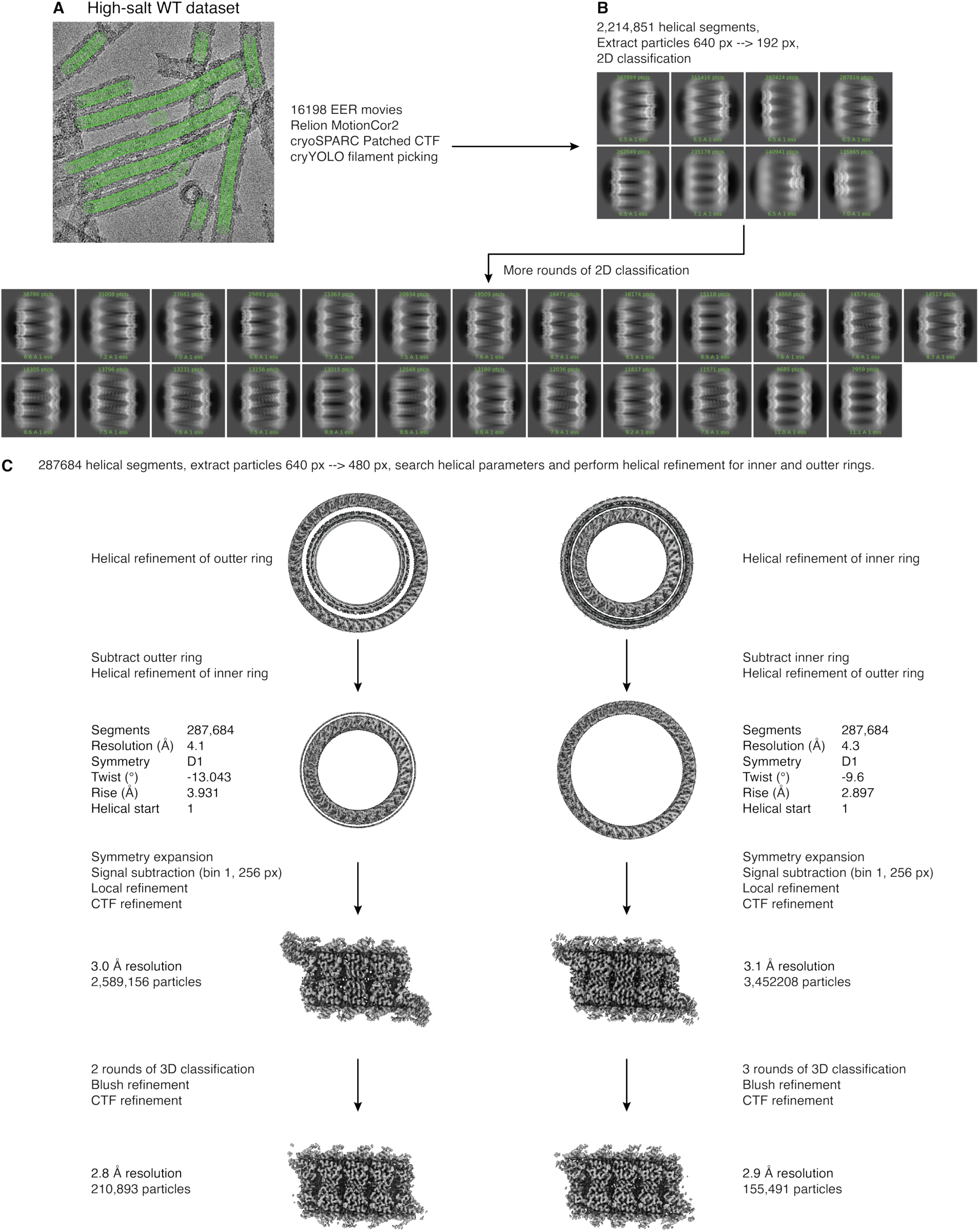
Image processing workflow for the in vitro wild-type (WT) M1 high-salt assembly. For a detailed description of the image processing, see materials and methods section. (**A**) Representative micrograph of the WT high-salt M1 assembly with picked filament segments indicated (green). (**B**) Representative 2D class averages of the WT high-salt assembly. Most classes contain two concentric M1 layers, whereas a subset contains three layers. Two-layer classes were selected for further cleaning by 2D classification. (**C**) For the selected particles, helical parameters were searched and helical refinements were performed separately for the inner and outer M1 layers. Reconstructed layers were subtracted to isolate the remaining layer for helical reconstruction. For each layer separately, particles were then processed by symmetry expansion, signal subtraction using a mask including 6 M1 monomers, and iterative local refinement and 3D classification for the inner and outer layers.

**Fig. S4.**
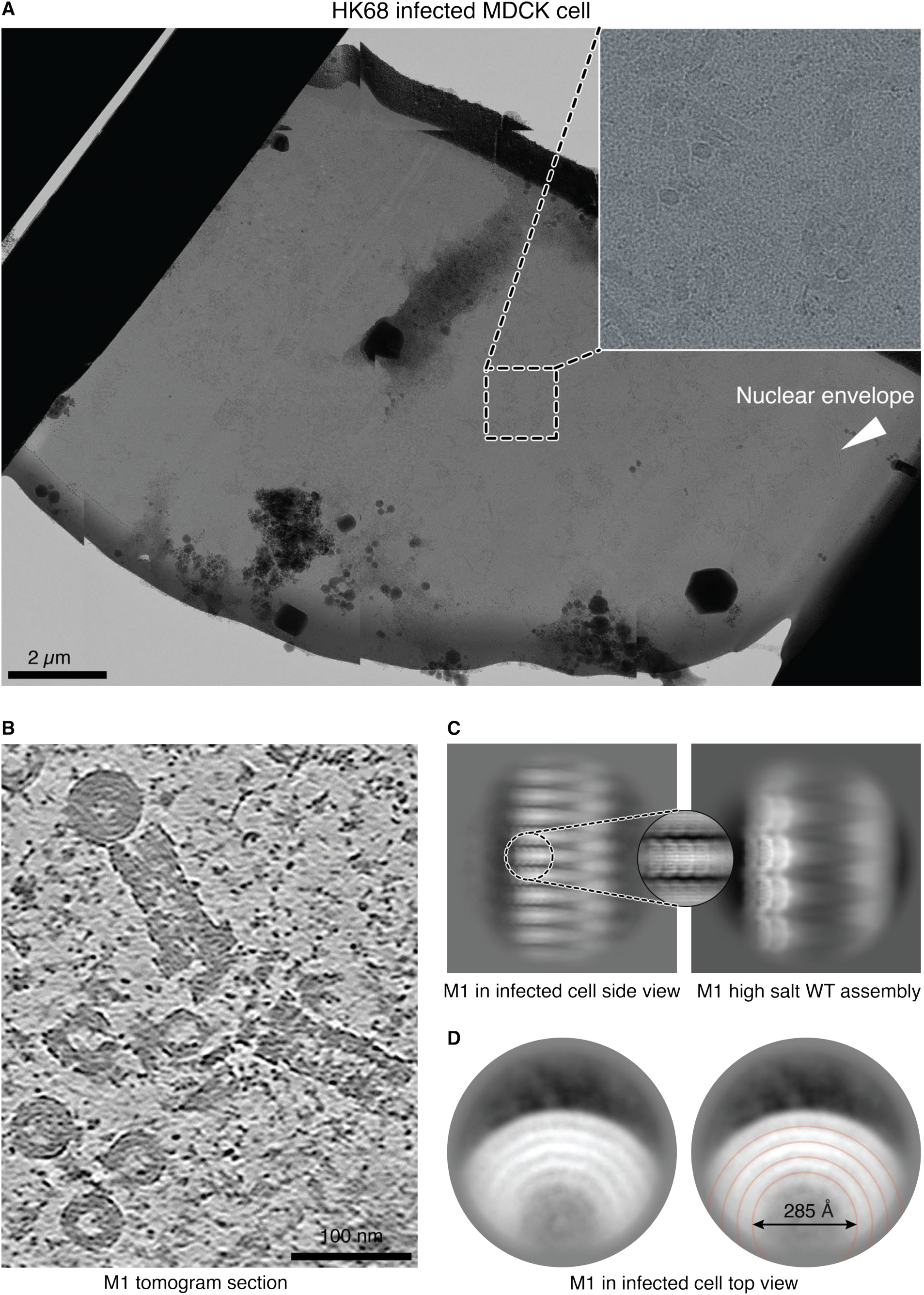
Cryo-FIB milling and analysis of M1 assemblies in HK68-infected cells. (**A**) Montage of a cryo-FIB–milled lamella from an HK68-infected MDCK cell. A zoomed view highlights multilayered M1 assemblies within the nucleus. (**B**) Representative tomographic slice showing multilayered M1 assemblies inside the nucleus. (**C**) Representative 2D class average of multilayered M1 assemblies in infected cells (left and inset) compared with a 2D class average of the three-layer WT high-salt M1 assembly (right). The overall packing is similar, although the cellular assemblies contain additional layers. (**D**) Representative 2D class average showing a top view of a five-layer M1 assembly in cells (left). Fitting concentric circles to the layers estimates the diameter of the innermost layer to be ∼285 Å, substantially narrower than most M1 filaments in virions (fig. S7C), indicating that M1_nucleotide_ can assemble into tightly curved M1 polymers similar to those seen at virion tips (Fig. 4 and figs. S13 and S14).

**Fig. S5.**
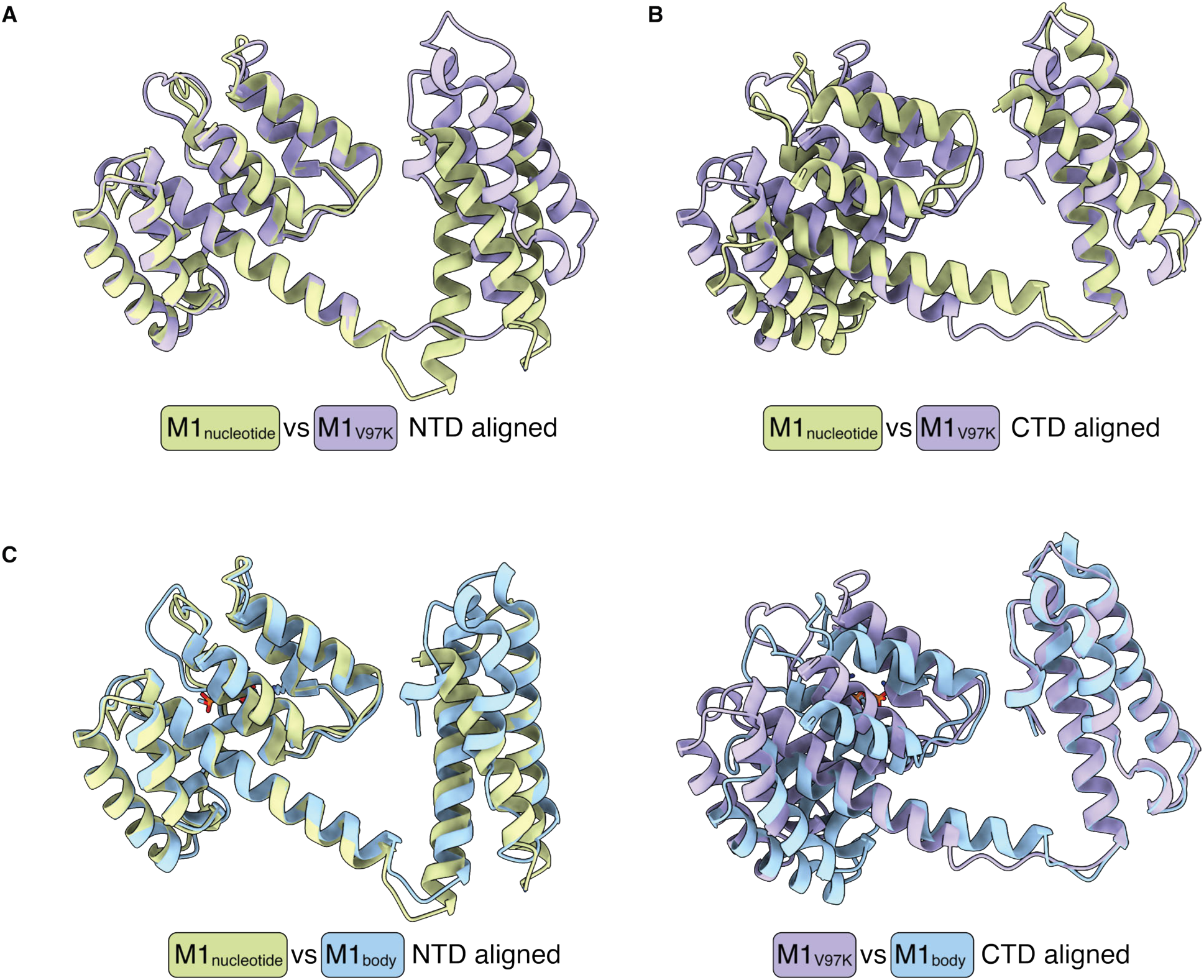
Comparisons of M1 conformations. (**A**) Overlay of M1_nucleotide_ (green) and M1_V97K_ (purple), aligned by the NTD. Switch 1 is partially helical in M1_nucleotide_ but is fully extended in M1_V97K_. (**B**) Overlay of M1_nucleotide_ (green) and M1_V97K_ (purple), aligned by the CTD. Switch 2 is associated with broad structural changes in the CTD: in M1_nucleotide_ the C-terminal residues extend the final alpha helix of M1 but these residues fold back to form a hooked configuration with a short 3_10_ helix in M1_V97K_. (**C**) Left, overlay of M1_nucleotide_ (green) and M1_body_ (blue); right, overlay of M1_V97K_ (purple) and M1_body_. In M1_body_, switch 1 is in the partially helical conformation seen in M1_nucleotide_, whereas switch 2 is in the hooked conformation seen in the M1_V97K_. M1_body_ therefore represents a hybrid of the two in vitro structures.

**Fig. S6.**
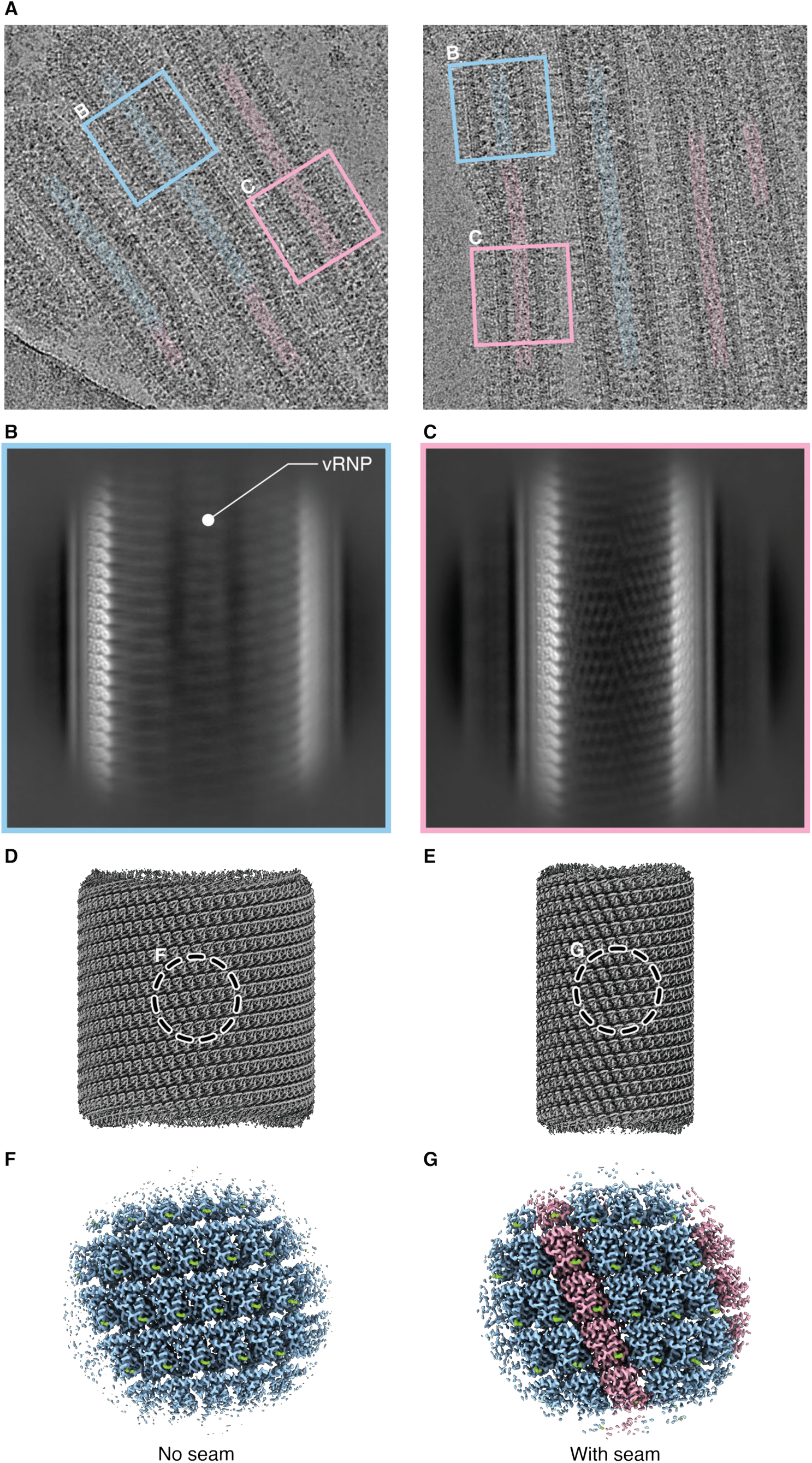
Wider and narrower regions of the HK68 virion body contain distinct M1 lattice organizations. (**A**) Representative micrographs of HK68 virions with the centers of picked filament segments indicated by colored circles. Segments were subjected to 2D classification as in fig. S7B, and segments assigned to wider and narrower classes are colored blue and pink, respectively. The wider segments typically localize to the front part of the virion body, whereas the narrower segments localize toward the rear. The wider segments accommodate the vRNPs and can extend beyond the vRNP-containing region (left). Blue and pink boxes indicate example particle segments contributing to the 2D classes shown in (B) and (C), respectively. (**B**) Representative 2D class average of the wider segments from fig. S7B. vRNP density is marked. (**C**) Representative 2D class average of the narrower segments from fig. S7B. (**D**) Representative helical reconstruction of the M1 lattice from the wider body region. Box indicates the region highlighted in (F). (**E**) Representative helical reconstruction of the M1 lattice from the narrower body region which corresponds to the bulk population. Box indicates the region highlighted in (G). (**F**) Asymmetric, high-resolution structure of a patch of the M1 filament from the wider region, as in fig. S7G. The structure is formed only from the M1_body_ conformation and does not contain a seam. (**G**) Asymmetric, high-resolution structure of a patch of the M1 filament from the narrower, bulk body population, as in Fig. 2E and fig. S7F. M1_seam_ is interspersed with M1_body_ at a 1:3 ratio.

**Fig. S7.**
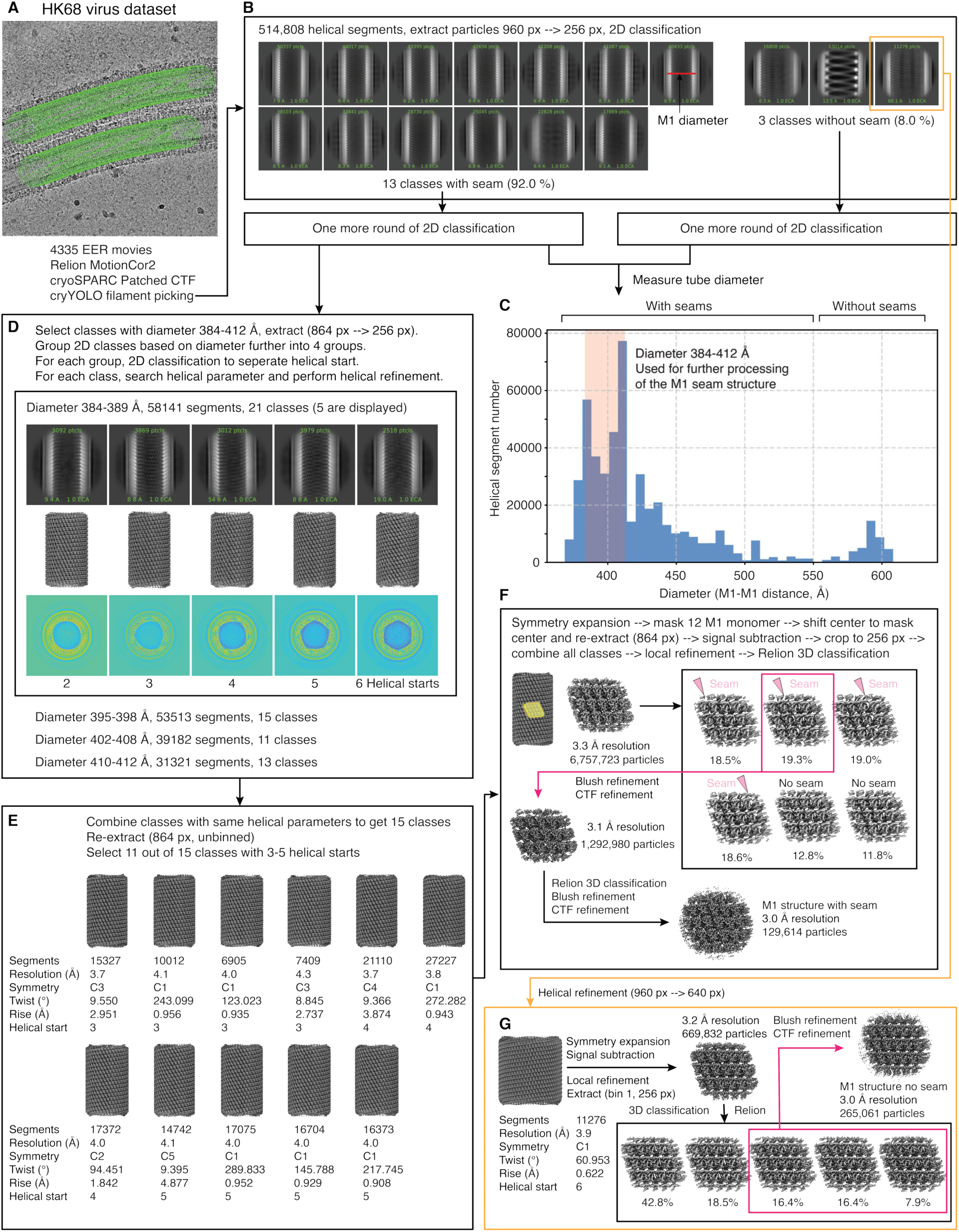
Image processing workflow for M1 structures from HK68 virions. For a detailed description of the image processing, see materials and methods section. (**A**) Representative micrograph of HK68 virions with picked filament segments indicated (green). (**B**) Representative 2D class averages of filaments with seam-containing M1 lattices (left) and of filaments without a seam (right). The red line in one class average indicates how the diameter of the M1 layer was measured. (**C**) Diameter distribution of M1 filaments. For seam-containing filaments, segments with diameters of 384–412 Å (pink shaded area) were selected for further processing. (**D**) Further 2D classification of diameter-matched, seam-containing filaments to separate subsets with different helical starts. Helical parameter searches and helical reconstructions were performed for all selected classes (60 reconstructions total). (**E**) Reconstructions with similar helical parameters were merged, yielding 15 higher-resolution reconstructions. 11 classes with three to five helical starts were selected for subsequent analyses. (**F**) For each selected class, a 12-subunit region of the M1 lattice was masked and the remaining signal was subtracted from particle images. Particles were re-extracted into 256-pixel boxes. Local refinement followed by 3D classification separated lattices with seams at distinct positions and seamless classes. Iterative Blush refinement and 3D classification produced the final reconstructions. (**G**) For seamless filaments, a single 2D class was selected and processed using the same masking, subtraction, refinement, and classification workflow to obtain the final reconstruction.

**Fig. S8.**
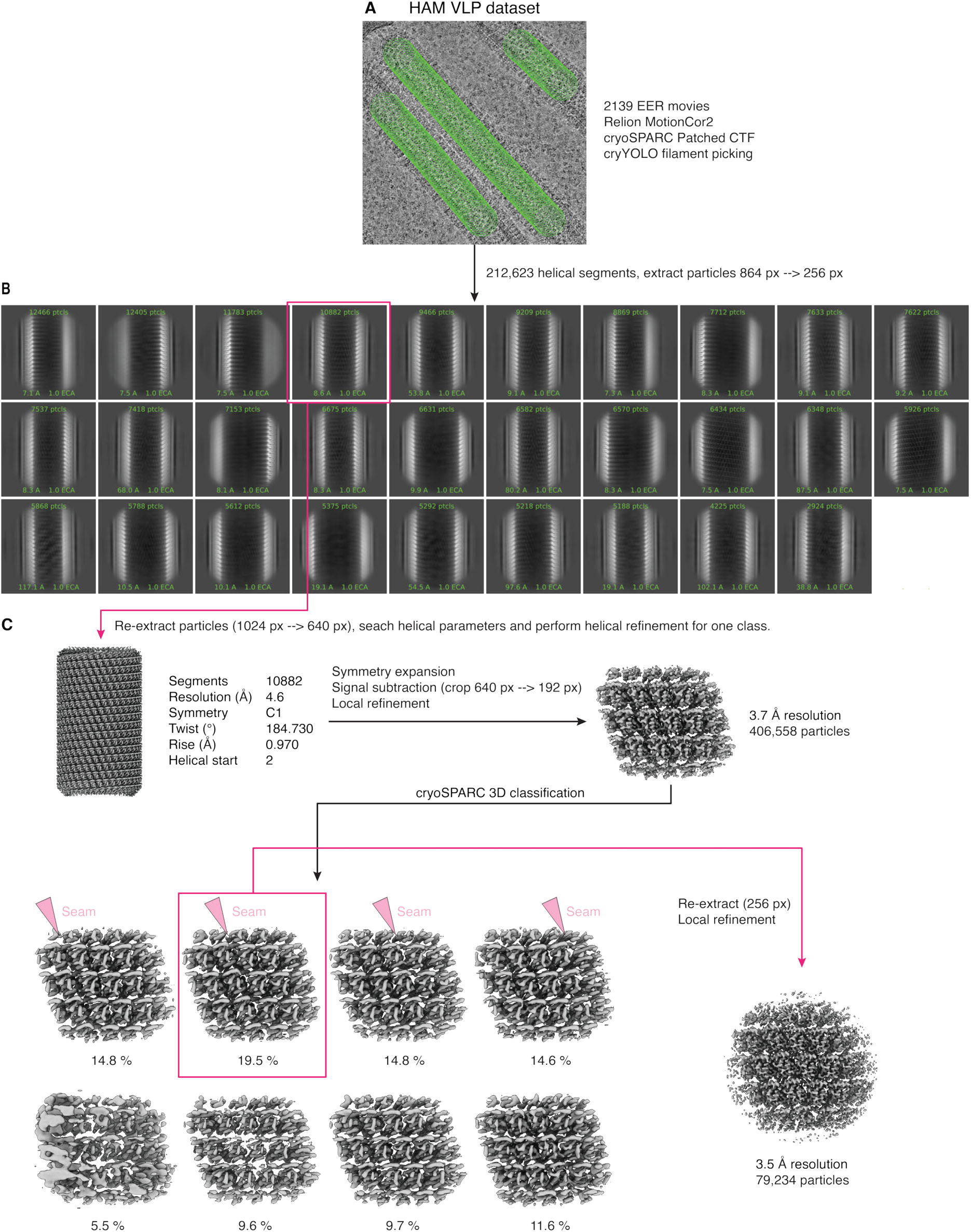
Image processing workflow for M1 structures from HAM VLPs. For a detailed description of the image processing, see materials and methods section. (**A**) Representative micrograph of HAM VLPs with picked filament segments indicated (green). (**B**) Representative 2D class averages of HAM VLP filaments. (**C**) A single 2D class was selected for helical parameter search and refinement. The final reconstruction was obtained using the same overall strategy as for HK68 virions.

**Fig. S9.**
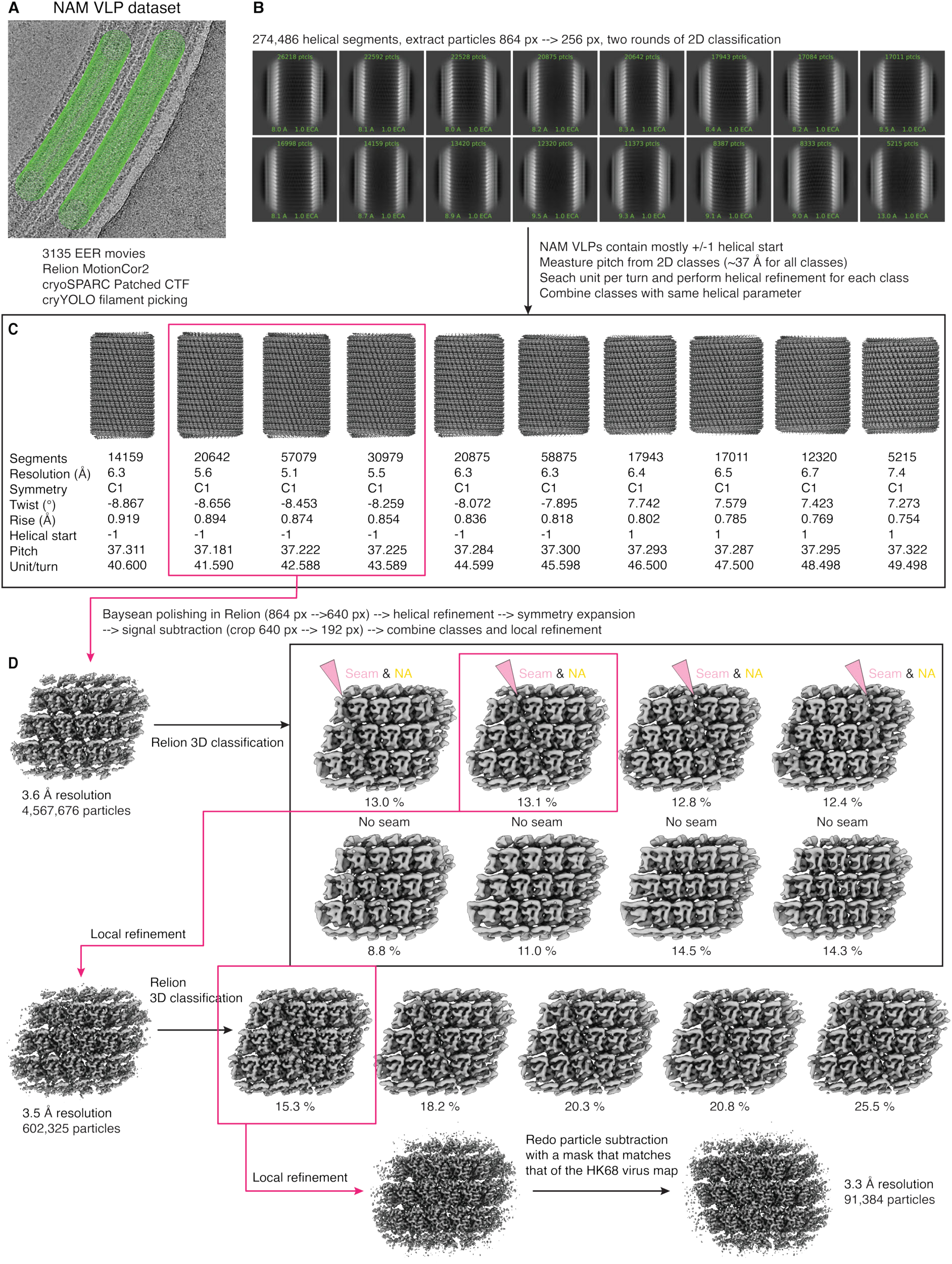
Image processing workflow for M1 structures from NAM VLPs. For a detailed description of the image processing, see materials and methods section. (**A**) Representative micrograph of NAM VLPs with picked filament segments indicated (green). (**B**) Representative 2D class averages of NAM VLP filaments. (**C**) Helical parameters were searched for individual 2D classes. Classes with similar parameters were merged to generate 10 initial 3D reconstructions. Three reconstructions with similar helical parameters were combined for further processing. (**D**) For each selected class, a 12-subunit region of the M1 lattice was masked and the remaining signal was subtracted from particle images. Particles were re-extracted into 256-pixel boxes. Local refinement followed by 3D classification separated lattices with seams at distinct positions and seamless classes. Iterative local refinement and 3D classification produced the final reconstructions. Signal subtraction was repeated using the HK68 12-subunit mask to enable direct map comparison.

**Fig. S10.**
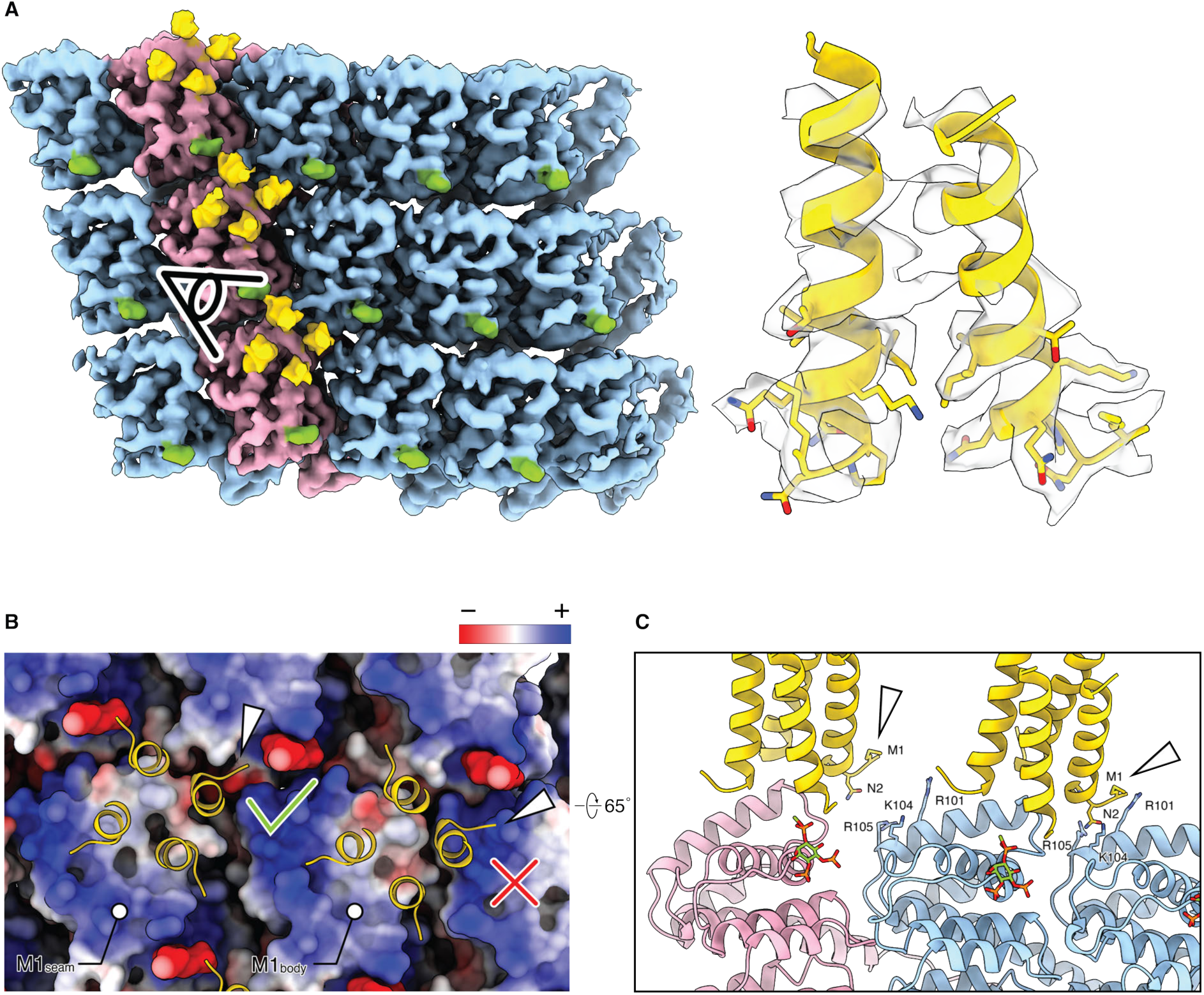
NA cytoplasmic-tail engagement at M1seam. (**A**) Left, viewing angle used for density visualization. Right, fit of the NA N-terminal model into the cryo-EM density. Density for several bulky side chains within the NA N-terminal residues is visible. (**B**) Comparison of NA engagement with M1_seam_ (left) and a hypothetical placement on M1_body_ (right). Electrostatic surface representation of the M1 lattice is shown. In the seam lattice, all four NA cytoplasmic tails occupy hydrophobic patches or intersubunit gaps. In the M1_body_ lattice, one NA tail would be positioned over a positively charged surface (white arrow, right), whereas the corresponding tail in the seam lattice sits in the groove between two M1 subunits (white arrow, left). (**C**) Rotated ribbon view of the model. Hypothetical placement of the NA cytoplasmic tail at the M1_body_ site produces clashes (white arrow, right), whereas the corresponding residues do not clash at the M1_seam_ binding site (white arrow, left).

**Fig. S11.**
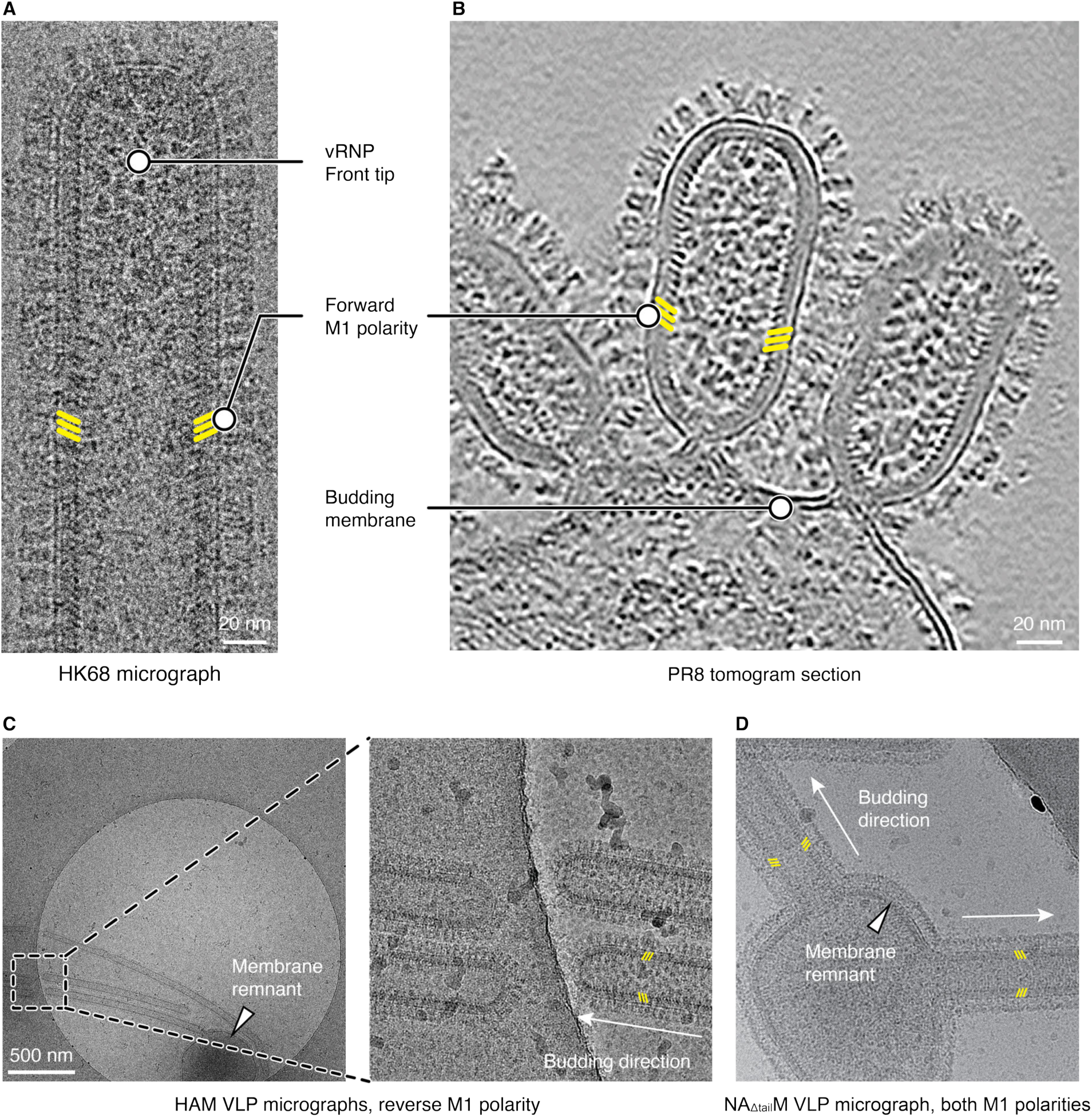
Polarity assignment for virions and VLPs. (**A**) Representative HK68 virion in a micrograph. The front tip was identified by the presence of clear vRNP density. M1 polarity was assigned from the direction of the CTD density (yellow markers). Forward polarity was assigned when the M1 CTD pointed away from the budding direction. HK68 virions show exclusively forward polarity. (**B**) Representative PR8 budding virion in a tomographic slice. Particles still connected to the plasma membrane were used to define the budding direction, and M1 polarity was assigned from CTD density orientation (yellow markers). PR8 virions show exclusively forward polarity. (**C**) HAM VLPs in micrographs. Left panel: a low-magnification micrograph was used to identify VLPs associated with a small membrane remnant (white arrowhead). The right panel shows a higher-magnification view of the HAM VLPs. Budding direction was defined as growth away from the membrane remnant (white arrowhead), and M1 polarity was assigned from CTD density orientation (yellow markers). In this example, the HAM VLP shows reverse polarity. whereas the two NA_Δtail_M VLPs show opposite polarities in the same micrograph. (**D**) Representative NA_Δtail_M VLPs at the same magnification as (C), right panel. As in (C), budding direction was defined as growth away from the membrane remnant (white arrowhead), and M1 polarity was assigned from CTD density orientation. In this example, the two NA_Δtail_M VLPs show opposite polarities.

**Fig. S12.**
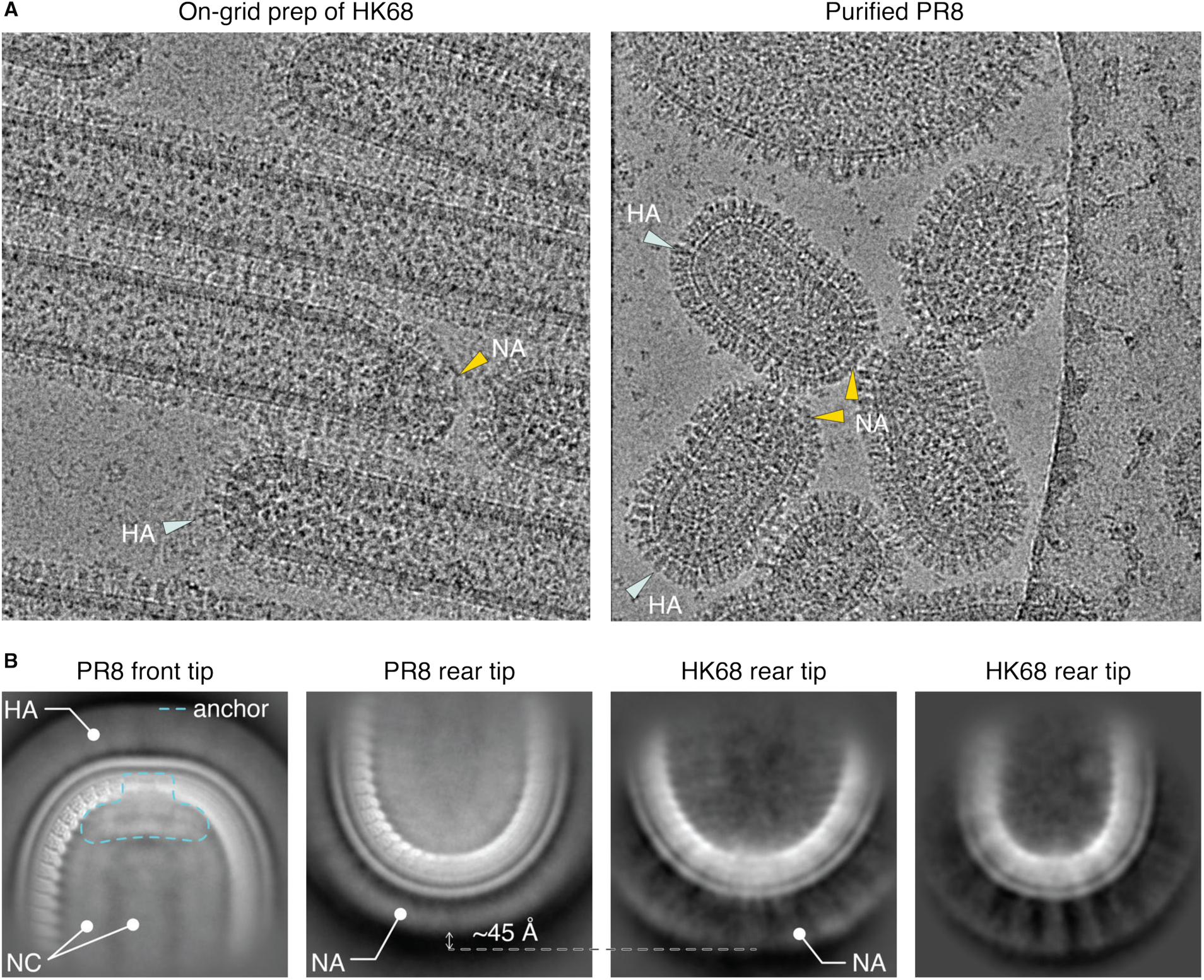
Tip organization of HK68 and PR8 virions. (**A**) Representative micrographs of on-grid–prepared HK68 virions (left) and purified PR8 virions (right). The front and rear tips show predominantly HA (mint arrows) and NA (gold arrows), respectively. (**B**) Representative 2D class averages of PR8 front and rear (same as Fig. 4D) tips, compared with two HK68 rear-tip class averages. The representative 2D class for HK68 front tip is shown in Fig. 4B. The PR8 front tip shows HA, nucleocapsid (NC), and anchor densities (cyan dashed outline), similar to those observed in HK68. HK68 NA appears 45 Å longer than PR8 NA, consistent with its extended hypervariable stalk region. Due to the smaller number of tip particle images and greater variability, HK68 rear-tip 2D classes were lower resolution.

**Fig. S13.**
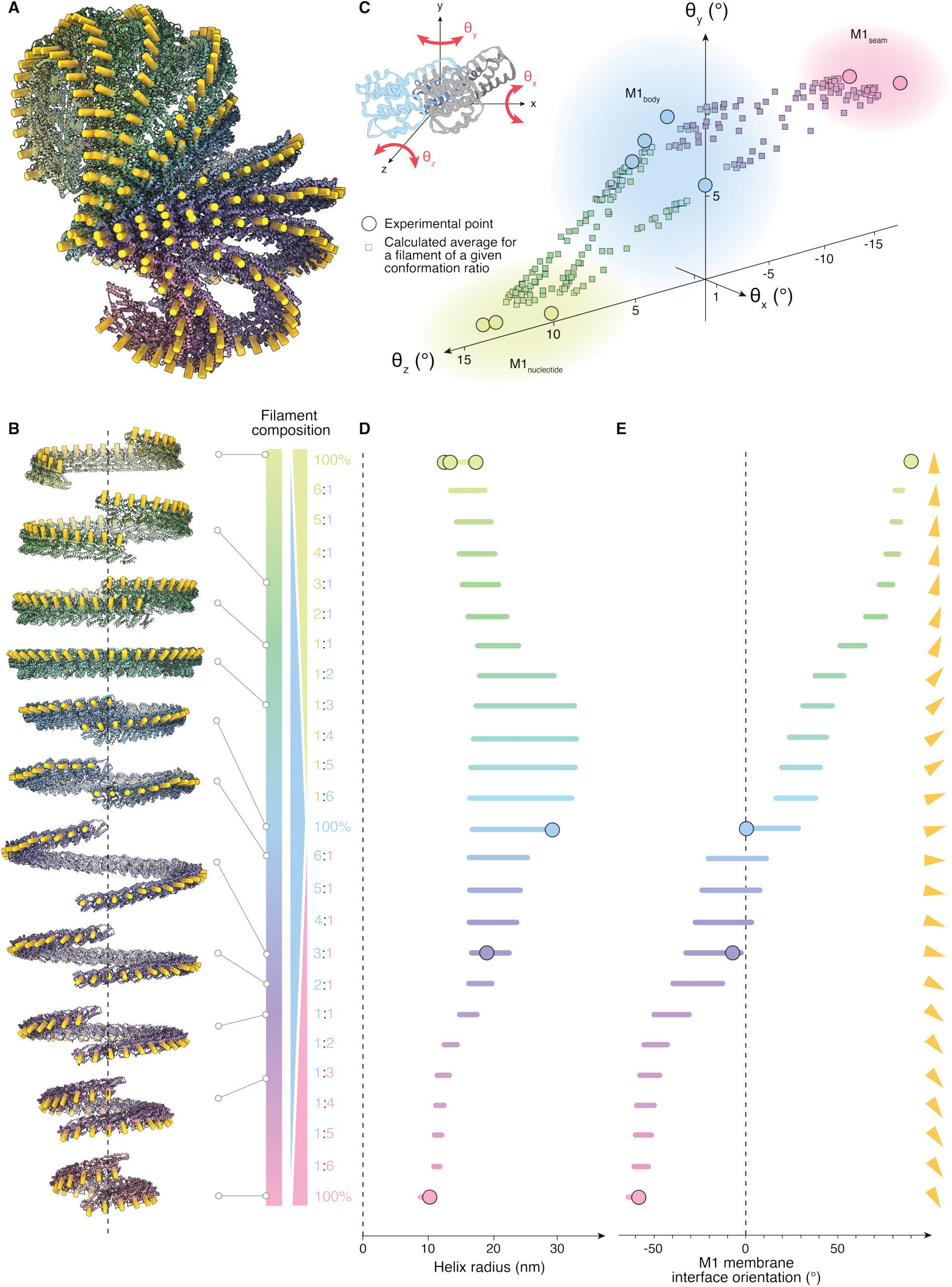
Modelled M1 polymer radius and membrane angle combining M1nucleotide, M1body and M1seam in different ratios. (**A**) Comparison of the propagation directions for modelled M1 polymers assembled with different ratios of M1_nucleotide_ (green), M1_body_ (blue) and M1_seam_ (pink), aligned relative to the NTD of the first monomer. Membrane-interaction surfaces are indicated by yellow bars. (**B**) The same M1 polymers aligned according to the filament axes (dotted line). Membrane-interaction surfaces are indicated by yellow bars. (**C**) The relative orientation of adjacent M1 monomers within the polymer can be described by three Euler angles (𝜃𝑥, 𝜃𝑦, 𝜃𝑧). The 3D plot shows the angles derived from experimentally determined structures (circles) and the average angles calculated for polymers built from different ratios of experimentally observed M1 conformations (squares). (**D**) Calculated radii across the ensemble of modelled M1 polymers. Values derived from experimentally determined structures are indicated by circles. (**E**) Calculated orientations of the membrane-interaction surface across the ensemble of modelled M1 polymers. Values derived from experimentally determined structures are indicated by circles. The orientation of the membrane-interaction surface is illustrated by yellow arrowheads. All panels are color-coded according to the composition of the M1 polymers.

**Fig. S14.**
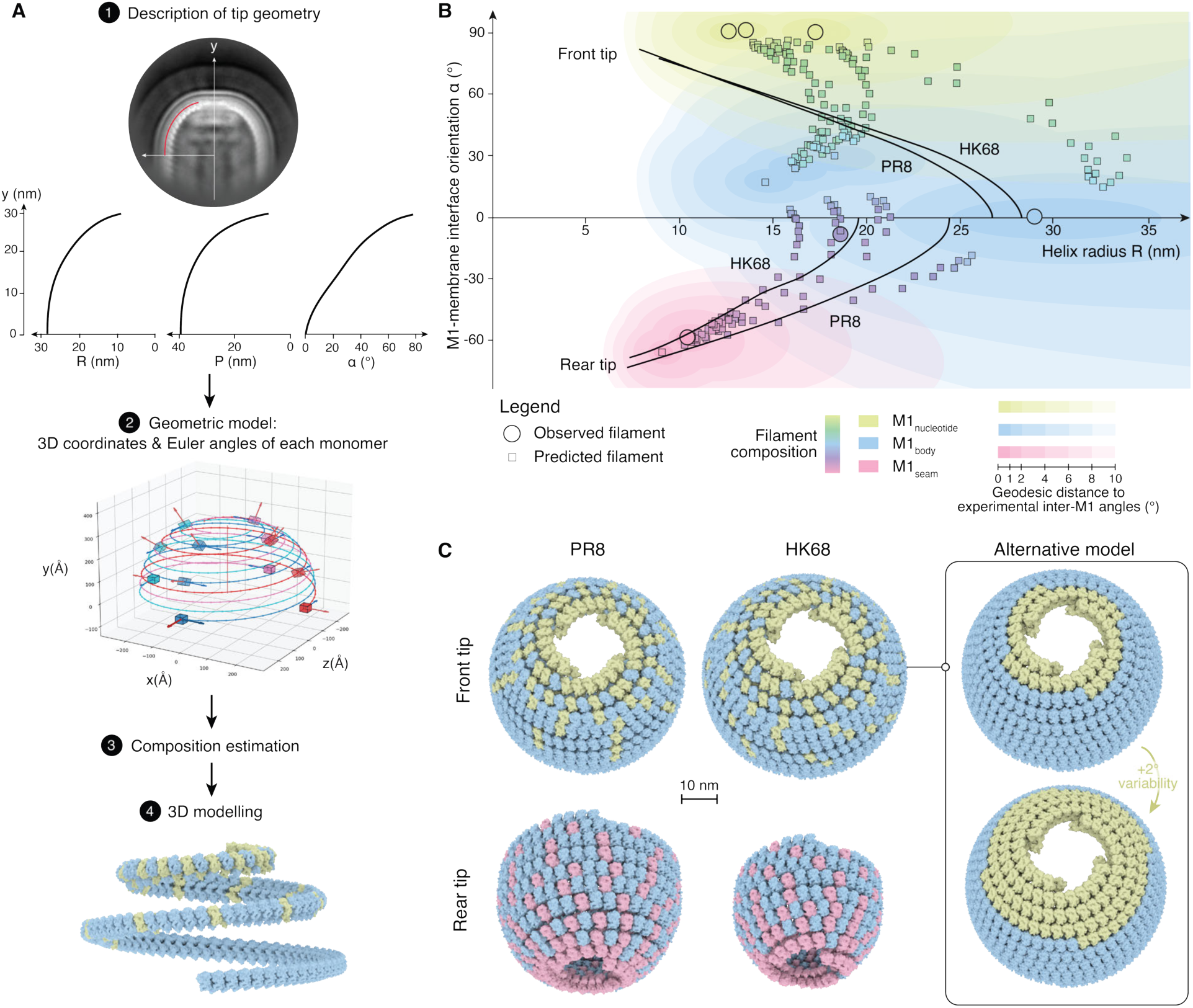
Modelling M1 polymers present at the front and rear tips of the virus. (**A**) Overview of the modelling workflow. Step 1: Tip geometry was described from 2D class averages of virion tips by marking a curve through the M1 NTD (red line), from which three continuous parameters were derived: radius 𝑅(𝑦), pitch 𝑃 (𝑦), and M1 membrane-binding-surface orientation 𝛼(𝑦). Step 2: Local 3D coordinates and pairwise Euler angles for each monomer were then obtained from the fitted functions 𝑅(𝑦), 𝑃 (𝑦) and 𝛼(𝑦), generating a geometric model. Step 3: Each monomer in the geometric model was assigned a conformation (M1_nucleotide_, M1_body_ or M1_seam_) according to the closest measured or predicted filament in Euler-angle space. Step 4: Molecular models were aligned to the reference frame, converted to polygonal meshes, and assembled on the resulting polymer geometry to generate 3D models of the virion tips. See methods for further details. (**B**) Comparison of predicted M1 polymer characteristics with those seen at the virion tips. The radius 𝑅(𝑦) and membrane-binding-surface orientation 𝛼(𝑦), as derived in (A), were plotted against one another (black lines) for representative front and rear tip classes of HK68 and PR8. Values of 𝑅 and 𝛼, calculated as in fig. 13, D and E, for experimentally determined helices are indicated by circles, and those for predicted mixed helices by squares. Gradually shaded areas indicate how allowing 0-10° deviations in pairwise Euler angles around each experimental value affects the corresponding polymer radius and membrane-binding-surface orientation. Such deviations can arise from inherent protein flexibility. (**C**) Predicted models of M1 filaments at the front (top) and rear (bottom) tips of PR8 (left) and HK68 (middle) virions, built from combinations of M1_nucleotide_ (green), M1_body_ (blue), and M1_seam_ (pink). Models were generated by gradually adjusting the conformation ratio to minimize local Euler angle deviations from the geometric model (left and middle). An alternative model for front tip formation is shown in which each monomer in the geometric model was assigned the geodesically closest experimental conformation without considering predicted mixed polymers. The upper model assumes no additional flexibility within the M1_nucleotide_ conformation, whereas the lower model allows an additional 2° of deviation, shifting the transition point to a different position along the polymer. See methods for further details.

**Fig. S15.**
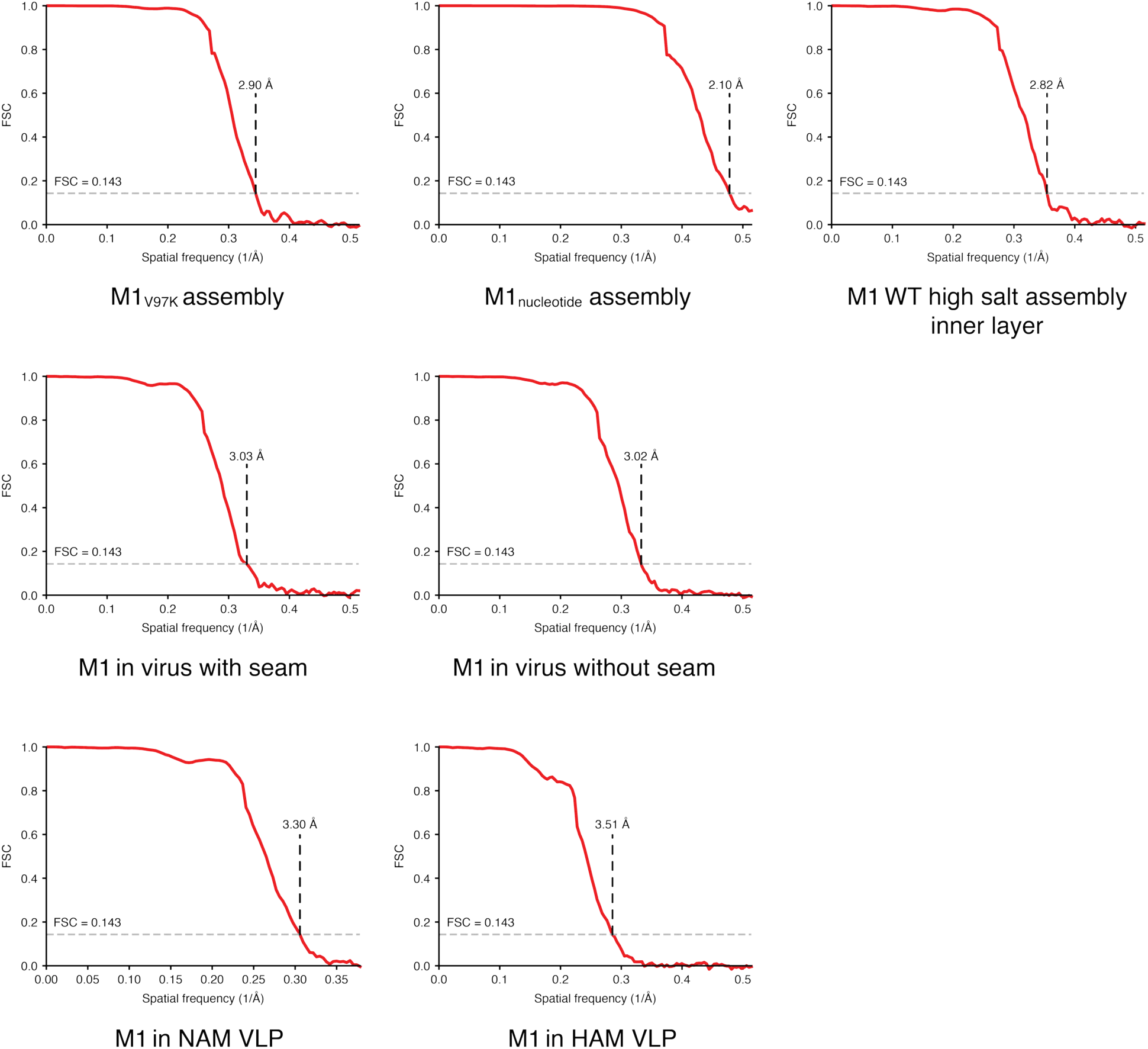
Gold-standard FSC curves for cryo-EM reconstructions.

**Table S1.** Cryo-EM data collection and atomic model refinement/validation statistics for M1 structures.

|  | M1<sub>v97K</sub><br>EMD: 57330<br>PDB: 29RJ | M1<sub>nucleotide</sub><br>EMD: 57331<br>PDB: 29RK | M1 WT<br>high salt<br>EMD: 57332<br>PDB: 29RL | M1 in HK68<br>virus with<br>seam<br>EMD: 57326<br>PDB: 29RG | M1 in HK68<br>virus without<br>seam<br>EMD: 57327<br>PDB: 29RH | M1 in NAM<br>VLP<br>EMD: 57328<br>PDB: 29RI | M1 in HAM<br>VLP<br>EMD: 57329 |
| --- | --- | --- | --- | --- | --- | --- | --- |
| **Data collection and processing** |  |  |  |  |  |  |  |
| Magnification | 130,000× | 130,000× | 130,000× | 130,000× | 130,000× | 130,000× | 130,000× |
| Voltage (kV) | 300 | 300 | 300 | 300 | 300 | 300 | 300 |
| Electron exposure (e<sup>-</sup>/Å<sup>2</sup>) | 50 | 44 | 40 | 45 | 45 | 43 | 48 |
| Defocus range (μm) | -1.0 to -3.0 | -1.0 to -2.5 | -1.0 to -3.0 | -1.2 to -3.2 | -1.2 to -3.2 | -1.5 to -3.5 | -1.0 to -3.0 |
| Pixel size (Å) | 0.9534 | 0.9534 | 0.9534 | 0.9534 | 0.9534 | 0.9534 | 0.9534 |
| Movies (no.) | 9,444 | 7,091 | 16,198 | 4,335 | 4,335 | 3,135 | 2,139 |
| Symmetry imposed | C1 | C1 | C1 | C1 | C1 | C1 | C1 |
| Initial particle images (segments) | 2,787,465 | 715,290 | 2,214,851 | 514,558 | 514,558 | 274,486 | 212,623 |
| Final particle images (no.) | 679,491 | 2,348,388 | 210,893 | 129,614 | 265,061 | 91,384 | 79,234 |
| Map resolution (Å) | 2.9 | 2.1 | 2.8 | 3.0 | 3.0 | 3.3 | 3.5 |
| FSC threshold | 0.143 | 0.143 | 0.143 | 0.143 | 0.143 | 0.143 | 0.143 |
| **Refinement** |  |  |  |  |  |  |  |
| Initial model used (PDB code) | 7JM3 | de novo | M1<sub>nucleotide</sub> | de novo | de novo | M1 in virus |  |
| Model resolution (Å) | 3.1 | 2.2 | 3.1 | 3.5 | 3.4 | 4.0 |  |
| FSC threshold | 0.5 | 0.5 | 0.5 | 0.5 | 0.5 | 0.5 |  |
| Map sharpening *B* factor (Å<sup>2</sup>) | -129.3 | -73.3 | -114.1 | -113.9 | -102.0 | -98.5 | -119.7 |
| **Model composition** |  |  |  |  |  |  |  |
| Non-hydrogen atoms | 1902 | 1892 | 1747 | 7832 | 1942 | 8242 |  |
| Protein residues | 251 | 244 | 235 | 1008 | 251 | 1084 |  |
| Ligands | 0 | 1 | 0 | 4 | 1 | 4 |  |
| ***B* factors (Å<sup>2</sup>)** |  |  |  |  |  |  |  |
| Protein | 61.66 | 30.99 | 52.80 | 98.15 | 81.30 | 97.09 |  |
| Ligand | - | 38.69 | - | 63.40 | 38.05 | 60.88 |  |
| **R.m.s. deviations** |  |  |  |  |  |  |  |
| Bond lengths (Å) | 0.003 | 0.008 | 0.003 | 0.003 | 0.003 | 0.003 |  |
| Bond angles (°) | 0.643 | 0.887 | 0.773 | 0.598 | 0.612 | 0.611 |  |
| **Validation** |  |  |  |  |  |  |  |
| MolProbity score | 1.08 | 0.75 | 0.69 | 0.68 | 1.04 | 0.73 |  |
| Clashscore | 2.87 | 0.79 | 0.58 | 0.51 | 2.57 | 0.73 |  |
| Poor rotamers (%) | 0 | 0 | 0 | 0 | 0 | 0 |  |
| **Ramachandran plot** |  |  |  |  |  |  |  |
| Favored (%) | 99.60 | 98.75 | 99.56 | 99.20 | 98.80 | 99.34 |  |
| Allowed (%) | 0.40 | 1.25 | 0.44 | 0.80 | 1.20 | 0.66 |  |
| Disallowed (%) | 0 | 0 | 0 | 0 | 0 | 0 |  |
| Q-score | 0.572 | 0.72 | 0.555 | 0.545 | 0.553 | 0.456 |  |

**Table S2.** Cryo-EM data collection and particle statistics for 2D class analyses.

|  | HK68 virus tips | PR8 virus tips | M1 in HK68 infected cells (lamellae) |
| --- | --- | --- | --- |
| **Data collection and processing** |  |  |  |
| Magnification | 130,000× | 130,000× | 105,000× |
| Voltage (kV) | 300 | 300 | 300 |
| Electron exposure (e<sup>-</sup>/Å<sup>2</sup>) | 45 | 40 | 53 |
| Defocus range (μm) | -1.2 to -3.2 | -1.0 to -3.0 | -2.5 to -4.5 |
| Pixel size (Å) | 0.9534 | 0.9534 | 1.2142 |
| Movies (no.) | 4,335 | 15,893 | 113 |
| Initial particle images (no.) | 4,106 | 14,284 | 7,669 segments (side)<br>7,538 (top) |
| Final particle images (no.) | 970 (front, Fig. 4B)<br>434 (rear, fig. S12B left)<br>386 (rear, fig. S12B right) | 1,768 (front, fig. S12B)<br>1,401 (rear, Fig. 4D and fig. S12B) | 2,229 (side, fig. S4C)<br>864 (top, fig. S4D) |

**Table S3.** Cryo-ET data collection parameters.

|  | PR8 virus (on-grid preparation) | M1 in HK68 infected cells (lamellae) |
| --- | --- | --- |
| **Data collection** |  |  |
| Magnification | 105,000× | 53,000× |
| Voltage (kV) | 300 | 300 |
| Electron exposure per tilt ($e^-/\text{\AA}^2$) | 3 | 3 |
| Defocus range ($\mu\text{m}$) | -2.5 to -5 | -2.5 to -5 |
| Pixel size ($\text{\AA}$) | 1.2142 | 2.3875 |
| Tilt series (no.) | 10 | 17 |
| Collection scheme | Dose-symmetric | Dose-symmetric |
| Tilt range ($^\circ$) | -60 to 60 | -60 to 60 |
| Tilt step ($^\circ$) | 3 | 3 |

## References

1. M. D. Badham, J. S. Rossman, Filamentous Influenza Viruses. Curr Clin Micro Rpt 3, 155–161 (2016).

2. B. Dadonaite, S. Vijayakrishnan, E. Fodor, D. Bhella, E. C. Hutchinson, Filamentous influenza viruses. Journal of General Virology 97, 1755–1764 (2016).

3. L. J. Calder, S. Wasilewski, J. A. Berriman, P. B. Rosenthal, Structural organization of a filamentous influenza A virus. Proceedings of the National Academy of Sciences 107, 10685–10690 (2010).

4. T. Noda, H. Sagara, A. Yen, A. Takada, H. Kida, R. H. Cheng, Y. Kawaoka, Architecture of ribonucleoprotein complexes in influenza A virus particles. Nature 439, 490–492 (2006).

5. A. J. Eisfeld, G. Neumann, Y. Kawaoka, At the centre: influenza A virus ribonucleoproteins. Nat Rev Microbiol 13, 28–41 (2015).

6. S. Wasilewski, L. J. Calder, T. Grant, P. B. Rosenthal, Distribution of surface glycoproteins on influenza A virus determined by electron cryotomography. Vaccine 30, 7368–7373 (2012).

7. J. J. Skehel, D. C. Wiley, Receptor Binding and Membrane Fusion in Virus Entry: The Influenza Hemagglutinin. Annu. Rev. Biochem. 69, 531–569 (2000).

8. J. Peukes, X. Xiong, S. Erlendsson, K. Qu, W. Wan, L. J. Calder, O. Schraidt, S. Kummer, S. M. V. Freund, H.-G. Kräusslich, J. A. G. Briggs, The native structure of the assembled matrix protein 1 of influenza A virus. Nature 587, 495–498 (2020).

9. E. R. Boulan, D. D. Sabatini, Asymmetric budding of viruses in epithelial monolayers: a model system for study of epithelial polarity. Proceedings of the National Academy of Sciences 75, 5071–5075 (1978).

10. P. Scheiffele, A. Rietveld, T. Wilk, K. Simons, Influenza Viruses Select Ordered Lipid Domains during Budding from the Plasma Membrane. Journal of Biological Chemistry 274, 2038–2044 (1999).

11. M. Takeda, G. P. Leser, C. J. Russell, R. A. Lamb, Influenza virus hemagglutinin concentrates in lipid raft microdomains for efficient viral fusion. Proceedings of the National Academy of Sciences 100, 14610–14617 (2003).

12. D. Dou, R. Revol, H. Östbye, H. Wang, R. Daniels, Influenza A Virus Cell Entry, Replication, Virion Assembly and Movement. Front. Immunol. 9, 1581 (2018).

13. M. J. Amorim, E. A. Bruce, E. K. C. Read, Á. Foeglein, R. Mahen, A. D. Stuart, P. Digard, A Rab11- and Microtubule-Dependent Mechanism for Cytoplasmic Transport of Influenza A Virus Viral RNA. Journal of Virology 85, 4143–4156 (2011).

14. A. J. Eisfeld, E. Kawakami, T. Watanabe, G. Neumann, Y. Kawaoka, RAB11A Is Essential for Transport of the Influenza Virus Genome to the Plasma Membrane. Journal of Virology 85, 6117–6126 (2011).

15. M. Enami, K. Enami, Influenza virus hemagglutinin and neuraminidase glycoproteins stimulate the membrane association of the matrix protein. Journal of Virology 70, 6653–6657 (1996).

16. A. Ali, R. T. Avalos, E. Ponimaskin, D. P. Nayak, Influenza Virus Assembly: Effect of Influenza Virus Glycoproteins on the Membrane Association of M1 Protein. Journal of Virology 74, 8709–8719 (2000).

17. D. Wang, A. Harmon, J. Jin, D. H. Francis, J. Christopher-Hennings, E. Nelson, R. C. Montelaro, F. Li, The Lack of an Inherent Membrane Targeting Signal Is Responsible for the Failure of the Matrix (M1) Protein of Influenza A Virus To Bud into Virus-Like Particles. Journal of Virology 84, 4673–4681 (2010).

18. A. Petrich, V. Dunsing, S. Bobone, S. Chiantia, Influenza A M2 recruits M1 to the plasma membrane: A fluorescence fluctuation microscopy study. Biophysical Journal 120, 5478–5490 (2021).

19. M. Hilsch, B. Goldenbogen, C. Sieben, C. T. Höfer, J. P. Rabe, E. Klipp, A. Herrmann, S. Chiantia, Influenza A Matrix Protein M1 Multimerizes upon Binding to Lipid Membranes. Biophysical Journal 107, 912–923 (2014).

20. S. Bobone, M. Hilsch, J. Storm, V. Dunsing, A. Herrmann, S. Chiantia, Phosphatidylserine Lateral Organization Influences the Interaction of Influenza Virus Matrix Protein 1 with Lipid Membranes. Journal of Virology 91, 10.1128/jvi.00267-17 (2017).

21. P. Raut, B. Obeng, H. Waters, J. Zimmerberg, J. A. Gosse, S. T. Hess, Phosphatidylinositol 4,5-Bisphosphate Mediates the Co-Distribution of Influenza A Hemagglutinin and Matrix Protein M1 at the Plasma Membrane. Viruses 14, 2509 (2022).

22. C. Chaimayo, T. Hayashi, A. Underwood, E. Hodges, T. Takimoto, Selective incorporation of vRNP into influenza A virions determined by its specific interaction with M1 protein. Virology 505, 23–32 (2017).

23. M. Wachsmuth-Melm, S. Peterl, A. O’Riain, J. Makroczyová, K. Fischer, T. Krischuns, S. Vale-Costa, M. J. Amorim, P. Chlanda, Visualizing influenza A virus assembly by in situ cryo-electron tomography. Nat Commun 16, 9394 (2025).

24. L. J. Mitnaul, M. R. Castrucci, K. G. Murti, Y. Kawaoka, The cytoplasmic tail of influenza A virus neuraminidase (NA) affects NA incorporation into virions, virion morphology, and virulence in mice but is not essential for virus replication. Journal of Virology 70, 873–879 (1996).

25. P. Chlanda, E. Mekhedov, H. Waters, A. Sodt, C. Schwartz, V. Nair, P. S. Blank, J. Zimmerberg, Palmitoylation Contributes to Membrane Curvature in Influenza A Virus Assembly and Hemagglutinin-Mediated Membrane Fusion. Journal of Virology 91, 10.1128/jvi.00947-17 (2017).

26. J. S. Rossman, R. A. Lamb, Influenza virus assembly and budding. Virology 411, 229–236 (2011).

27. K. Martin, A. Heleniust, Nuclear transport of influenza virus ribonucleoproteins: The viral matrix protein (M1) promotes export and inhibits import. Cell 67, 117–130 (1991).

28. K. Martin, A. Helenius, Transport of incoming influenza virus nucleocapsids into the nucleus. Journal of Virology 65, 232–244 (1991).

29. B. Sha, M. Luo, Structure of a bifunctional membrane-RNA binding protein, influenza virus matrix protein M1. Nat Struct Mol Biol 4, 239–244 (1997).

30. S. Arzt, F. Baudin, A. Barge, P. Timmins, W. P. Burmeister, R. W. H. Ruigrok, Combined Results from Solution Studies on Intact Influenza Virus M1 Protein and from a New Crystal Form of Its N-Terminal Domain Show That M1 Is an Elongated Monomer. Virology 279, 439–446 (2001).

31. L. Selzer, Z. Su, G. D. Pintilie, W. Chiu, K. Kirkegaard, Full-length three-dimensional structure of the influenza A virus M1 protein and its organization into a matrix layer. PLOS Biology 18, e3000827 (2020).

32. O. Terrier, V. Moules, C. Carron, G. Cartet, E. Frobert, M. Yver, A. Traversier, T. Wolff, B. Riteau, N. Naffakh, B. Lina, J.-J. Diaz, M. Rosa-Calatrava, The influenza fingerprints: NS1 and M1 proteins contribute to specific host cell ultrastructure signatures upon infection by different influenza A viruses. Virology 432, 204–218 (2012).

33. F. J. Milder, M. Jongeneelen, T. Ritschel, P. Bouchier, I. J. M. Bisschop, M. de Man, D. Veldman, L. Le, B. Kaufmann, M. J. G. Bakkers, J. Juraszek, B. Brandenburg, J. P. M. Langedijk, Universal stabilization of the influenza hemagglutinin by structure-based redesign of the pH switch regions. Proceedings of the National Academy of Sciences 119, e2115379119 (2022).

34. C. M. Mair, T. Meyer, K. Schneider, Q. Huang, M. Veit, A. Herrmann, A Histidine Residue of the Influenza Virus Hemagglutinin Controls the pH Dependence of the Conformational Change Mediating Membrane Fusion. Journal of Virology 88, 13189–13200 (2014).

35. R. Kanai, K. Kar, K. Anthony, L. H. Gould, M. Ledizet, E. Fikrig, W. A. Marasco, R. A. Koski, Y. Modis, Crystal Structure of West Nile Virus Envelope Glycoprotein Reveals Viral Surface Epitopes. Journal of Virology 80, 11000–11008 (2006).

36. P. Várnai, T. Balla, Visualization of Phosphoinositides That Bind Pleckstrin Homology Domains: Calcium- and Agonist-induced Dynamic Changes and Relationship to Myo-[3H]inositol-labeled Phosphoinositide Pools. J Cell Biol 143, 501–510 (1998).

37. S. L. Winter, G. Golani, F. Lolicato, M. Vallbracht, K. Thiyagarajah, S. S. Ahmed, C. Lüchtenborg, O. T. Fackler, B. Brügger, T. Hoenen, W. Nickel, U. S. Schwarz, P. Chlanda, The Ebola virus VP40 matrix layer undergoes endosomal disassembly essential for membrane fusion. EMBO J 42, EMBJ2023113578 (2023).

38. C. E. Mire, D. Dube, S. E. Delos, J. M. White, M. A. Whitt, Glycoprotein-Dependent Acidification of Vesicular Stomatitis Virus Enhances Release of Matrix Protein. Journal of Virology 83, 12139–12150 (2009).

39. J. Peukes, S. Dmitrieff, F. J. Nédélec, J. A. G. Briggs, A physical model for M1-mediated influenza A virus assembly. Biophysical Journal 124, 134–144 (2025).

40. J. Blok, G. M. Air, Variation in the membrane-insertion and “stalk” sequences in eight subtypes of influenza type A virus neuraminidase. Biochemistry 21, 4001–4007 (1982).

41. A. M. Ernst, S. Zacherl, A. Herrmann, M. Hacke, W. Nickel, F. T. Wieland, B. Brügger, Differential transport of Influenza A neuraminidase signal anchor peptides to the plasma membrane. FEBS Letters 587, 1411–1417 (2013).

42. D. N. Mastronarde, Automated electron microscope tomography using robust prediction of specimen movements. Journal of Structural Biology 152, 36–51 (2005).

43. H. Guo, E. Franken, Y. Deng, S. Benlekbir, G. S. Lezcano, B. Janssen, L. Yu, Z. A. Ripstein, Y. Z. Tan, J. L. Rubinstein, Electron-event representation data enable efficient cryoEM file storage with full preservation of spatial and temporal resolution. IUCrJ 7, 860–869 (2020).

44. A. Punjani, J. L. Rubinstein, D. J. Fleet, M. A. Brubaker, cryoSPARC: algorithms for rapid unsupervised cryo-EM structure determination. Nat Methods 14, 290–296 (2017).

45. T. Wagner, F. Merino, M. Stabrin, T. Moriya, C. Antoni, A. Apelbaum, P. Hagel, O. Sitsel, T. Raisch, D. Prumbaum, D. Quentin, D. Roderer, S. Tacke, B. Siebolds, E. Schubert, T. R. Shaikh, P. Lill, C. Gatsogiannis, S. Raunser, SPHIRE-crYOLO is a fast and accurate fully automated particle picker for cryo-EM. Commun Biol 2, 218 (2019).

46. S. H. W. Scheres, RELION: Implementation of a Bayesian approach to cryo-EM structure determination. Journal of Structural Biology 180, 519–530 (2012).

47. D. Kimanius, K. Jamali, M. E. Wilkinson, S. Lövestam, V. Velazhahan, T. Nakane, S. H. W. Scheres, Data-driven regularization lowers the size barrier of cryo-EM structure determination. Nat Methods 21, 1216–1221 (2024).

48. F. W. Studier, Protein production by auto-induction in high-density shaking cultures. Protein Expression and Purification 41, 207–234 (2005).

49. K. Jamali, L. Käll, R. Zhang, A. Brown, D. Kimanius, S. H. W. Scheres, Automated model building and protein identification in cryo-EM maps. Nature 628, 450–457 (2024).

50. P. Emsley, K. Cowtan, Coot: Model-building tools for molecular graphics. Acta Crystallographica Section D: Biological Crystallography 60, 2126–2132 (2004).

51. T. I. Croll, ISOLDE: A physically realistic environment for model building into low-resolution electron-density maps. Acta Crystallographica Section D: Structural Biology 74, 519–530 (2018).

52. P. D. Adams, P. V. Afonine, G. Bunkóczi, V. B. Chen, I. W. Davis, N. Echols, J. J. Headd, L. W. Hung, G. J. Kapral, R. W. Grosse-Kunstleve, A. J. McCoy, N. W. Moriarty, R. Oeffner, R. J. Read, D. C. Richardson, J. S. Richardson, T. C. Terwilliger, P. H. Zwart, PHENIX: A comprehensive Python-based system for macromolecular structure solution. Acta Crystallographica Section D: Biological Crystallography 66, 213–221 (2010).

53. T. D. Goddard, C. C. Huang, E. C. Meng, E. F. Pettersen, G. S. Couch, J. H. Morris, T. E. Ferrin, UCSF ChimeraX: Meeting modern challenges in visualization and analysis. Protein Science 27, 14–25 (2018).

